# Biochemical dissection of a fungal highly reducing polyketide synthase condensing region reveals basis for acyl group selection

**DOI:** 10.1101/2024.12.23.630095

**Authors:** Mia E. Foran, Nazia B. Auckloo, Y. T. Candace Ho, Shaonan Liu, Yang Hai, Matthew Jenner

## Abstract

Fungal highly reducing polyketide synthases (hrPKSs) are remarkable multidomain enzymes that catalyse the biosynthesis of a diverse range of structurally complex compounds. During biosynthesis, the ketosynthase (KS) and acyltransferase (AT) domains of the condensing region are visited by the acyl carrier protein (ACP) domain during every cycle, catalysing chain priming and elongation reactions. Despite their significance, our comprehension of how these steps contribute to biosynthetic fidelity remains poorly understood. Herein, we report the full *in vitro* reconstitution of each catalytic step carried out by a hrPKS condensing region. Application of intact protein mass spectrometry shows that the AT domain operates selective acyl transfer to the ACP domain, ensuring loading of the correct starter / extender units, which is governed by a substrate recycling mechanism and kinetic parameters towards each substrate. The KS domain is shown to select for the correct starter unit for transacylation and subsequent chain elongation, yet harbours flexibility towards the extender unit supplied. This work provides the first comprehensive interrogation of a fungal hrPKS condensing region, which both affirms and expands our understanding of how these systems function, and provides a framework for future bioengineering efforts.

## INTRODUCTION

Fungal iterative polyketide synthases (iPKSs) are responsible for the biosynthesis of many structurally diverse and biologically significant polyketide natural products.^1,2^ These remarkable multi-domain enzymes are categorised as non-reducing (nrPKS), partially reducing (prPKS), or highly reducing (hrPKS), according to their reductive processing capability.^3–5^ Of these, hrPKSs closely resemble the mammalian fatty acid synthase (mFAS) in both domain architecture and catalysis.^6^ In these megasynthases, an iterative cycle of decarboxylative Claisen condensation reactions between malonyl extender units bound to the acyl carrier protein (ACP) domain and the growing acyl chain is catalysed by the ketosynthase (KS) domain; with extender units continually loaded onto the ACP domain by an acyl transferase (AT) domain. Intermediates and extender units are covalently tethered via a thioester bond to a 4’-phosphopantetheine (Ppant) prosthetic ‘arm’ that is post-translationally appended to the ACP domain.^7,8^

During fatty acid biosynthesis, the β-keto group is reduced after each chain extension cycle to a methylene via the sequential action of ketoreductase (KR), dehydratase (DH) and enoylreductase (ER) domains. However, hrPKSs introduce structural complexity by programming the degree of β-carbon processing, allowing installation of β-hydroxyl groups (KR alone), α, β−double bonds (KR and DH), and methylene groups (KR, DH and ER). Furthermore, some hrPKSs harbour an active methyltransferase (MT) domain allowing methylation of the α-carbon; a reaction that has been shown to play an important role in programming.^9,10^ It should be noted that mFAS contains a catalytically inactive methyltransferase-like (MT^0^) domain, preventing this modification and resulting in the production of unbranched fatty acids.^11^

From both a structural and biosynthetic perspective, hrPKSs can be split into two functional sections: i). the condensing region, comprised of the KS and AT domains bridged by a structural linker domain (LD), required for starter unit selection and chain extension reactions, and ii). the modifying region, consisting of MT, KR, DH and ER domains that are used in various combinations during each chain extension cycle (**Fig. 1a**).^6,12^ A combination of *in vivo* and *in vitro* studies have provided valuable insights into the functioning of hrPKS domains, indicating that kinetic competition for the ACP-tethered intermediates by the catalytic domains appears to underpin biosynthetic programming.^9,10,13–20^ However, to date, much of this work has focussed on domains from the modifying region, which are visited in various combinations during product biosynthesis. In contrast, domains of the condensing region are visited during every catalytic cycle, and likely play an important role in ensuring biosynthetic fidelity of the product via tight control over starter / extender unit recruitment (AT domain), and selection / extension of the correct biosynthetic intermediate (KS domain). Both of these domains catalyse two-step reactions, which have proved difficult to fully unpick using traditional methods (e.g. radiolabelling assays / small molecule substrate mimics), and as a result are understudied for their roles in hrPKS programming.

**Figure 1.**
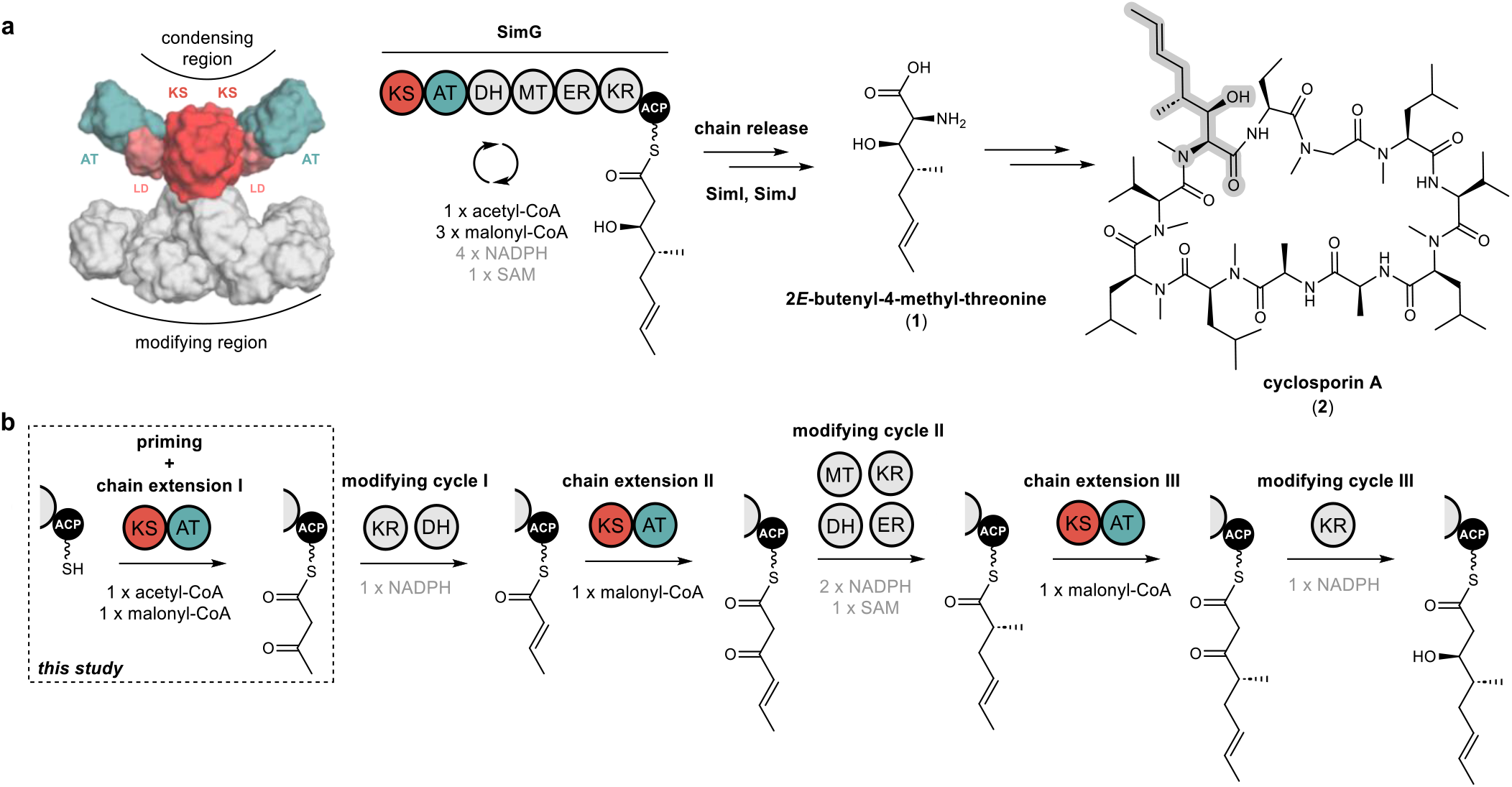
Biosynthesis of 2*E*-butenyl-4-methyl threonine (Bmt) by the SimG hrPKS. **a**). Domain arrangement of the SimG hrPKS responsible for the biosynthesis of the polyketide-derived amino acid *E*-2-butenyl-4-methyl-threonine (Bmt) incorporated into cyclosporin A (highlighted in grey). The catalytic KS and AT domains that comprise the condensing region are highlighted in red and teal, respectively, and the structural LD is highlighted in pink. The structural model of SimG was constructed using AlphaFold. **b**). Schematic showing each chain extension and modifying cycle during the biosynthesis of the polyketide Bmt-precursor. The catalytic domains, substrates and co-factors required for each step are shown.

In this work we have functionally dissected the condensing region from the SimG hrPKS, responsible for producing the polyketide scaffold of the non-proteinogenic amino acid (2*E*)-butenyl-4-methyl-L-threonine (Bmt, **1**) (**Fig. 1a**).^21^ This unit is incorporated into the immunosuppressant cyclosporin A (**2**), for which the Bmt side chain provides important contacts with its molecular target, cyclophilin A.^22,23^. Previous work suggests that SimG selects acetyl-CoA as a starter unit, followed by three chain extension reactions with malonyl-CoA to yield a tetraketide product, with each catalytic cycle utilising different combinations of modifying domains (**Fig. 1b**).^24–26^ The short biosynthetic trajectory of this hrPKS makes it an excellent model system to study the intrinsic molecular programming of these intriguing megaenzymes. Here, using a combination of excised catalytic domains and intact protein mass spectrometry (MS), we report the full *in vitro* reconstitution of each individual catalytic step carried out by the SimG hrPKS condensing region. Our results demonstrate that the substrate loading in hrPKSs is a tightly controlled process, with biosynthetic fidelity ensured by intrinsic substrate selectivity of each catalytic domain.

## RESULTS

### Heterologous production of 3*R*-hydroxyl-4*R*-methyl-6*E*-octenoic acid by SimG hrPKS

Previous studies have shown that SimG can produce the precursor to Bmt (**1**), 3*R*-hydroxyl-4*R*-methyl-6*E*-octenoic acid (Hma, **3**), both *in vivo* and *in vitro*.^25–27^ Building on these observations, the *simG* gene was cloned from a cDNA library of *Tolypocladium inflatum* NRRL 8044 and expressed under the control of ADH2p promoter in *Saccharomyces cerevisiae* JHY686 as a heterologous host (**Supplementary Table S1**).^28,29^ Following a 3-day fermentation, analysis of the culture supernatant by LC-ESI-MS showed production of Hma (**3**), and subsequent purification allowed structure confirmation by NMR spectroscopy (**Fig. 2, Supplementary Table S2**).

**Figure 2.**
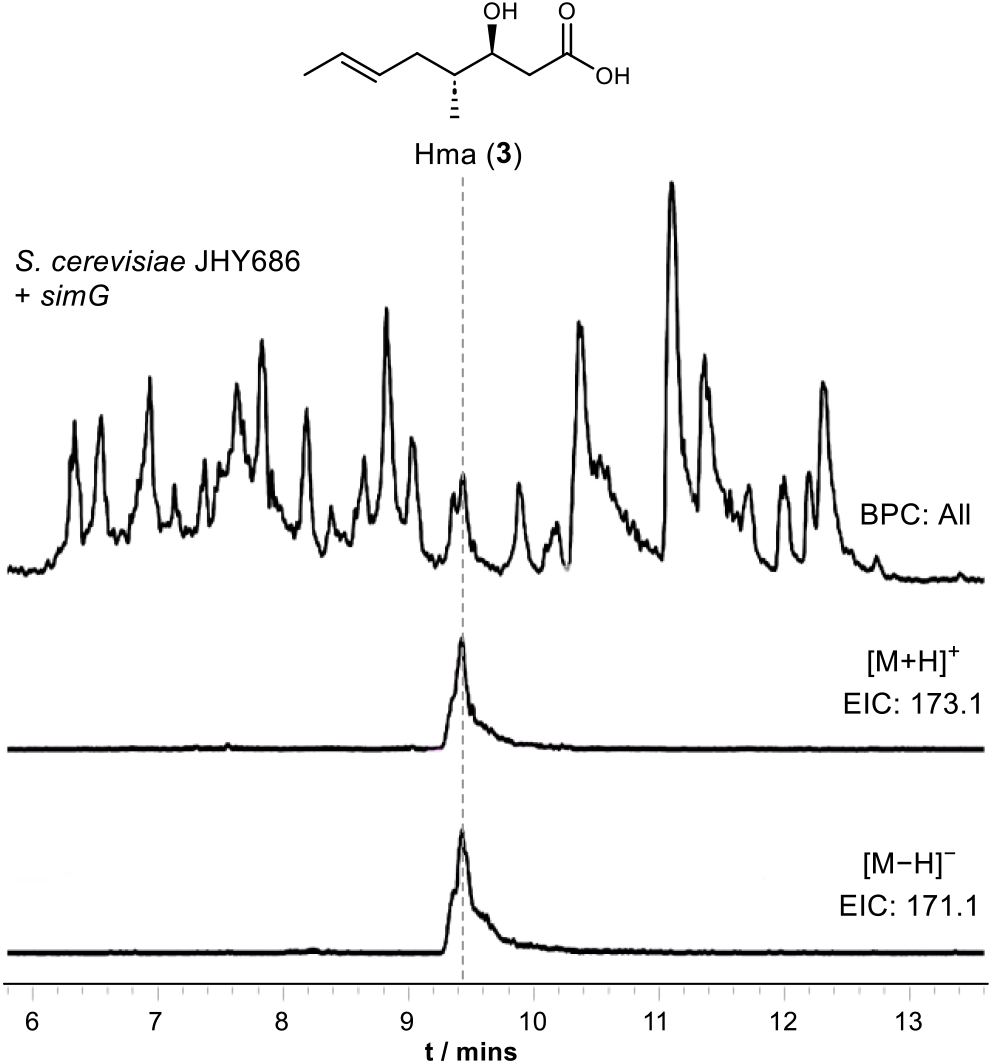
Detection of 3*R*-hydroxyl-4*R*-methyl-6*E*-octenoic acid (Hma) from *S. cerevisiae* JHY686. LC-ESI-MS base peak chromatogram (top) of ethyl acetate extract from culture medium of *S. cerevisiae* JHY686 expressing the *simG* gene. Extracted ion chromatograms for [M+H]^+^, *m/z* = 173.1 (*middle*) and [M−H]^−^, *m/z* = 171.1 (*bottom*) corresponding to Hma (**3**). Purification followed by NMR spectroscopy verified the structure of the product (**Supplementary Table S2**).

### Dissection of the condensing region and ACP domain

Having established that SimG biosynthesises the expected product, we proceeded to construct expression plasmids that contained excised domains from the condensing region to allow KS and AT domains to be assayed with the ACP domain (**Supplementary Table S3**). Here, domain boundaries were identified through sequence and structural alignment with LovB, for which a high-resolution structure has been reported,^12^ and the condensing region has been previously dissected.^15^ The ACP domain produced soluble protein and could be converted to its *holo*-form using the phosphopantetheinyl transferase, Sfp,^30^ whilst the excised AT domain required the flanking LD region to generate a soluble protein (**Supplementary Fig. S1a** and **b**).

It was not possible to produce a soluble construct of the KS domain alone, however, the KS-AT didomain could be produced at good levels. Despite this, following Ni^2+^-NTA purification the protein rapidly degraded into fragments at ∼60 kDa and ∼45 kDa, suggesting that the protein was proteolytically degraded into KS and AT portions, likely cutting within the extensive (65 amino acid) disordered linker between the KS and AT domains that contains several cleavage sites for *E. coli* proteases (**Supplementary Fig. S1c** and **d**). This was also observed for the LovB KS-AT didomain, but anion exchange chromatography prevented degradation.^15^ The SimG KS-AT didomain degraded within hours of Ni^2+^-NTA purification, preventing downstream chromatography, and addition of protease inhibitors during purification did not alleviate this issue. Comparison of the equivalent disordered linker in LovB revealed that it is significantly shorter (20 amino acids) than that of SimG. In an effort to produce an intact KS-AT didomain, we excised the middle segment, making it a similar length to the LovB linker (**Supplementary Fig. S1c**). The resulting SimG KS-AT(Δ474-515) didomain construct produced soluble protein which did not degrade (**Supplementary Fig. S1e**), and using this construct as a template, a KS-AT^0^ (S705A) mutant was also produced to allow the KS to be assayed in isolation (**Supplementary Fig. S1e**), however a KS^0^-AT^0^ (C215A, S705A) double mutant yielded insoluble protein.

### Starter and extender unit selection catalysed by the AT domain

Previous work has demonstrated that fungal hrPKS AT domains possess dual-specificity, loading both starter and extender units in a similar manner to mFAS.^15^ Here, the starter unit is typically acetyl, although some notable exceptions exist including propionyl (e.g. pseurotin A),^31^ benzoyl (e.g. strobilurin and squalestatin S1),^32,33^ and nicotinyl (e.g. pyripyropene),^34^ whilst the extender unit is exclusively malonyl. The AT domain catalyses transfer of acyl units from the corresponding CoA to the Ppant of the ACP domain in a two-step process.^35^ Firstly, the active site serine residue (Ser705 in SimG) is acylated by the corresponding acyl-CoA. This is followed by transfer of the acyl group to the Ppant thiol of the ACP domain, with both steps being inherently reversible (**Fig. 3a**).

**Figure 3.**
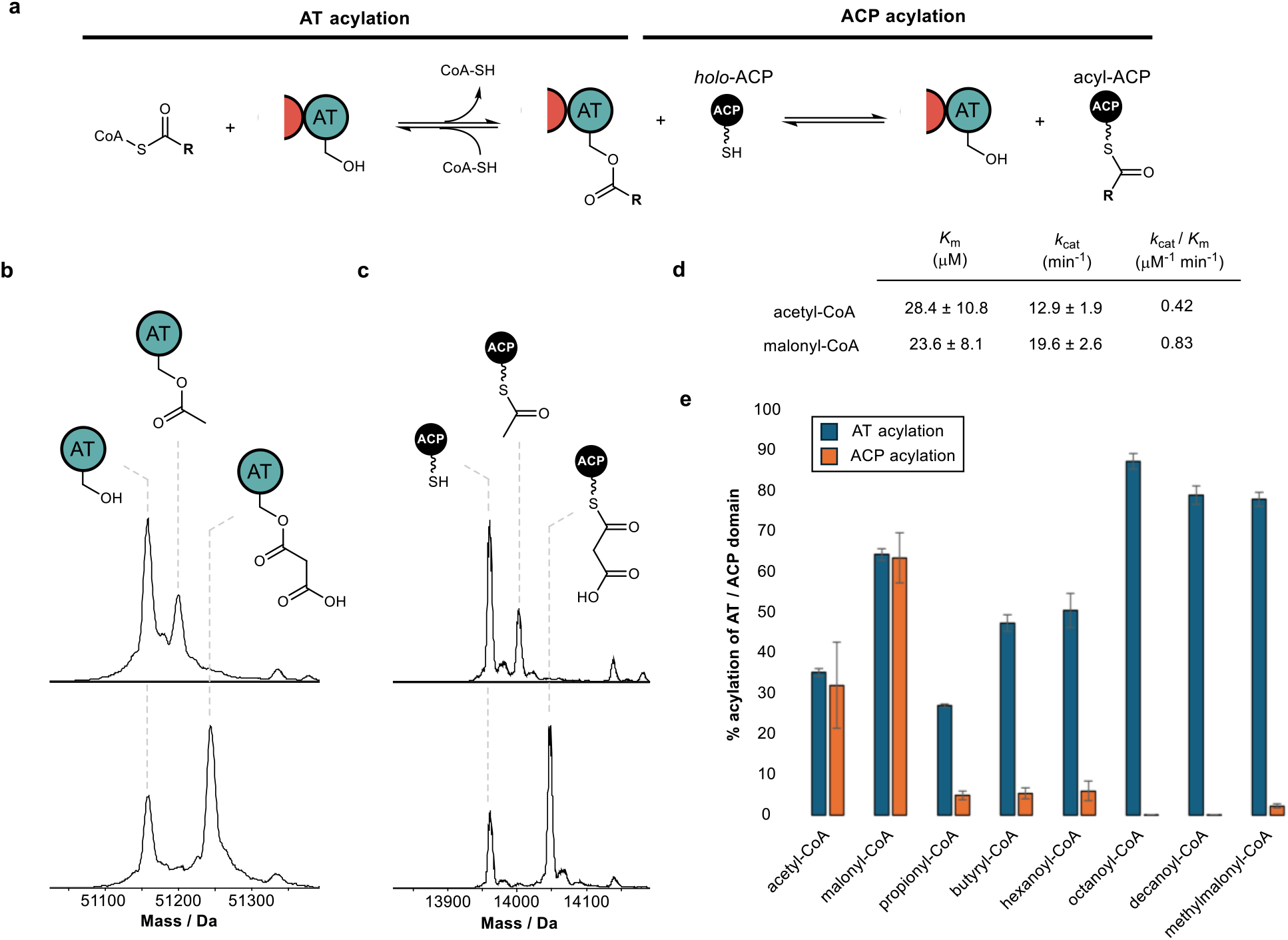
Starter and extender unit selectivity of the SimG AT domain. **a**). Schematic of AT-catalysed loading of acyl groups onto the ACP domain from coenzyme A thioesters. **b**). Deconvoluted mass spectra of SimG AT domain following incubation with acetyl-CoA (*top*) and malonyl-CoA (*bottom*), and **c**). deconvoluted mass spectra of SimG ACP domain after incubation with SimG AT and acetyl-CoA (*top*) and malonyl-CoA (*bottom*). Experiments were conducted in triplicate and representative spectra are shown. **d**). Michaelis–Menten properties for the rate of ACP domain acylation catalysed by the SimG AT domain for acetyl- and malonyl-CoA. The SimG AT domain exhibits a 2-fold preference (k_cat_ / K_m_) for transacylating malonyl-versus acetyl-CoA. **e**). Bar chart showing the percentage acylation of SimG AT domain (*blue*), and subsequent acyl transfer to the ACP domain (*orange*), following incubation with a range of acyl-CoAs. Error bars represent the standard deviation of three replicates.

The biosynthesis of Hma (**3**) requires one acetyl-CoA and three malonyl-CoA units, in line with previous labelling studies,^24^ to achieve the tetraketide product. Using our purified SimG AT domain, we first assayed the ability of acetyl-CoA and malonyl-CoA to transacylate the AT domain using intact protein mass spectrometry.

Incubation of SimG AT with an excess of acetyl or malonyl-CoA yielded 35.3 ± 0.9 % loading of acetyl and 64.5 ± 1.4 % loading of malonyl, indicated by mass shifts of +42 Da and +86 Da, respectively (**Fig. 3b**). This preference towards malonyl has been observed for the LovB AT domain,^15^ and is likely due to the electrostatic interaction between a highly conserved arginine residue in the AT domain active site (R730 in SimG AT) and the carboxylate anion of malonyl unit – an interaction not possible for acetyl.^36^ Inclusion of the *holo*-ACP domain resulted in equivalent levels of acyl transfer to the Ppant thiol, loading 32.1 ± 10.6 % acetyl and 63.7 ± 6.2 % malonyl (**Fig. 3c**), with no transfer observed when the AT domain was omitted (**Supplementary Fig. S2**). The same assay allowed us to determine the steady-state kinetic parameters for the system by quantifying acyl-ACP formation over time (**Fig. 3d** and **Supplementary Fig. S3**). Whilst the SimG AT domain exhibits broadly similar kinetic constants to the LovB AT domain^15^, the measured *k*_cat_ value for acetyl transfer was much higher, resulting in a 2-fold preference for transacylating malonyl-versus acetyl-CoA (*k*_cat_ / *K*_m_: acetyl = 0.42 uM^-1^ min^-1^, malonyl = 0.83 uM^-1^ min^-1^). This is in stark contrast to a 20-fold preference observed for the LovB AT domain, and may indicate that AT domains have tuned their processivity according to the number of malonyl-CoA extender units required to assemble the final product (i.e. 8 malonyl-CoA for LovB, 3 malonyl-CoA for SimG). It is worth acknowledging that the kinetic data obtained here, as in other studies, uses dissected domains. This is in contrast to the covalent tethering within the native hrPKS, meaning that local concentration of the ACP domain is high, promoting short diffusion pathways.^37^

We next sought to explore whether alternative starter and extender units could be accepted by the AT domain and transferred to the ACP domain. First, a range of acyl-CoAs with increasing saturated carbon chain lengths were tested. Whilst good levels of AT transacylation were observed for C3 – C10 substrates, only trace levels of transfer to the ACP domain were detected, with octanoyl and decanoyl units not transferred at all (**Fig. 3e**). We next tested whether methylmalonyl-CoA could be used as an alternative extender unit by the AT domain. Similar to the trend observed for non-native starter units, methylmalonyl-CoA was able to acylate the AT domain, but could only transfer trace amounts to the ACP domain. The ability of the AT domain to become acylated by non-native starter / extender units, but not catalyse subsequent transfer to the ACP domain efficiently, suggests that once attached to the active site Ser705 residue, the non-native acyl chains are unable to find the correct catalytic geometry for the second step of the reaction. However, acylation of the AT domain by a non-native starter / extender unit from the cellular environment would, in principle, inhibit the hrPKS without a mechanism for removal (*vide infra*). These observations suggest that although some non-native starter / extender units might be able to prime the ACP domain at low levels, it seems unlikely that they would provide the required substrate flux to assemble the polyketide product at scalable quantities. Furthermore, the intrinsic selectivity of the AT domain for acetyl- and malonyl-CoA means that attempts to produce derivatives via alternative starter / extender units would need to be conducted *in vitro* where the substrates can be controlled, or with entire AT domain swapping.

### The AT domain mediates starter / extender unit recycling

Our data has demonstrated that, similar to mFAS, the SimG AT domain exhibits dual-specificity towards acetyl and malonyl units. In a cellular context, the same AT domain must therefore select from a substrate pool containing both acetyl- and malonyl-CoA, which will act as competitive inhibitors of each other. To overcome this problem, the mFAS AT domain operates a substrate sorting mechanism, rapidly exchanging acetyl and malonyl units between CoA and the ACP Ppant thiol. Only when the correct acyl unit is loaded does a productive reaction occur, either transfer to the KS active site cysteine for acetyl, or chain extension for malonyl. At a given timepoint, if the incorrect substrate is loaded, the substrate is transferred back to CoA via the AT domain, making the system dependent upon the availability of free CoA.^38^ Whilst a similar mechanism is assumed for fungal hrPKSs based on their similarities to mFAS, this has yet to be investigated.

To examine whether this phenomenon occurs in hrPKSs, *apo*-SimG ACP was converted to its acetyl- and malonyl-ACP forms using Sfp and the corresponding acyl-CoA (**Fig. 4**, trace i). Incubation with CoA-SH or the AT domain alone showed no removal of the acetyl or malonyl groups compared to the control (**Fig. 4**, traces ii and iii). However, the combination of the AT domain with excess CoA-SH resulted in removal of both acetyl (31.1 ± 7.5 %) and malonyl (41.4 ± 3.2 %) units, with the malonyl unit appearing to be more readily removed than acetyl (**Fig. 4**, trace iv). The propensity of acetyl units to be more resistant to offloading may allow the required ‘dwell time’ for subsequent transfer to the KS domain, whilst the more labile malonyl unit provides opportunities for the AT domain to select an acetyl starter unit from the cellular pool, if required. Indeed, a time-course of our original acyl loading experiments (*vida supra*) with acetyl- and malonyl-CoA in competition resulted in malonyl preferentially loaded during the early time points, before an accumulation of acetyl units becomes the major species later in the assay (**Supplementary Fig. S4**). This pattern is reflective of the relative loading and offloading efficiencies of each acyl unit and demonstrates that AT domains from hrPKSs adopt a substrate sorting mechanism, similar to that of mFAS, in order to select the correct acyl unit from the cellular pool.

**Figure 4.**
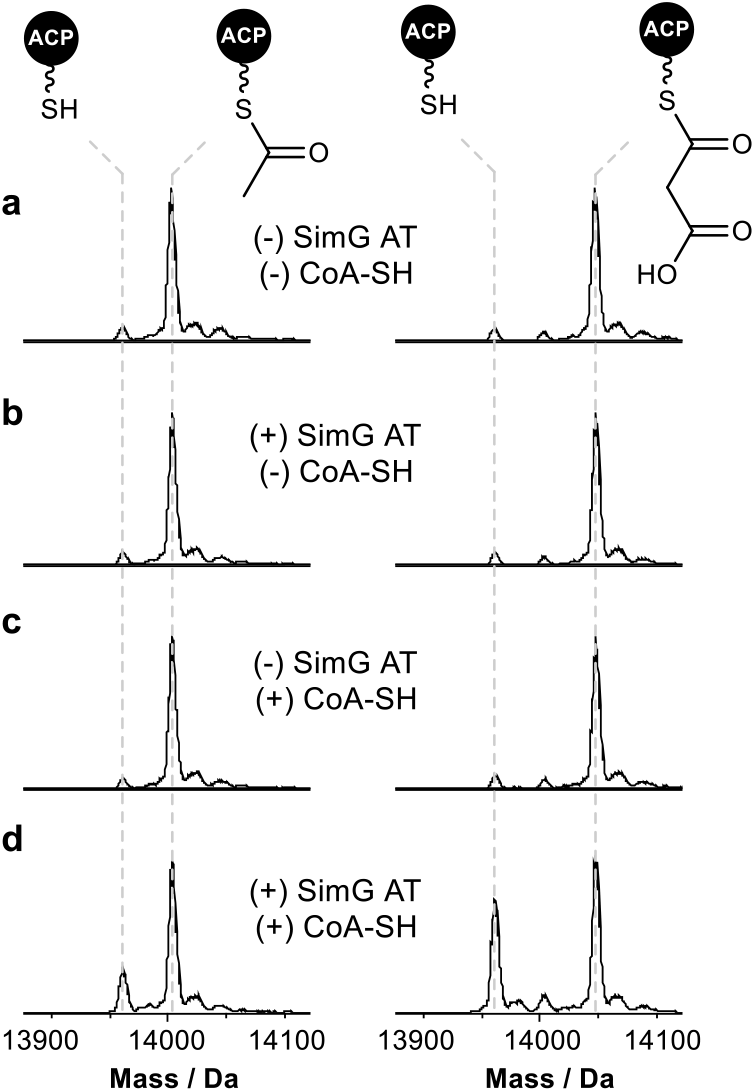
Starter / extender unit recycling catalysed by the SimG AT domain. **a**). Deconvoluted mass spectra of SimG acetyl-ACP (*left*) and malonyl-ACP (*right*), and following incubation with **b**). SimG AT alone, **c**). CoA-SH alone, and **d**). SimG AT and CoA-SH. Experiments were conducted in triplicate and representative spectra are shown.

Whilst the non-native starter / extender units tested previously were not efficiently transferred to the ACP domain, all were able to acylate the AT domain to good levels (**Fig. 3e**). Following pre-acylation with each non-native starter / extender unit, incubation of excess CoA-SH resulted in offloading of propionyl, butyryl, hexanoyl,

octanoyl and decanoyl units with a decreasing efficiency with respect to chain length, and low-levels of offloading observed for the methylmalonyl species (**Supplementary Fig. S5**). The ability to remove these aberrant acyl groups from the AT domain using free CoA is an essential feature of the enzyme to ensure it does not become inhibited by other acyl-CoAs within the cellular environment.

### The KS domain exhibits selectivity for the correct biosynthetic intermediate

Having established that the SimG AT domain selectively loads acetyl and malonyl units onto the ACP domain, we next turned our attention to the KS domain. The catalytic activity of the KS domain can be broken down into two steps – i). transacylation of the KS active site Cys residue with the correct biosynthetic intermediate, and ii). chain extension via a Clasien-like condensation reaction between the acyl-KS and malonyl-ACP, yielding a β-keto thioester product (**Fig. 5a**). Whilst the selectivity of the starter and extender units is tightly governed by the AT domain, whether the KS domain is able to select for the required ACP-bound intermediate with respect to chain-length, methylation, or extent of β-keto processing remains unclear, and to date untested.

**Figure 5.**
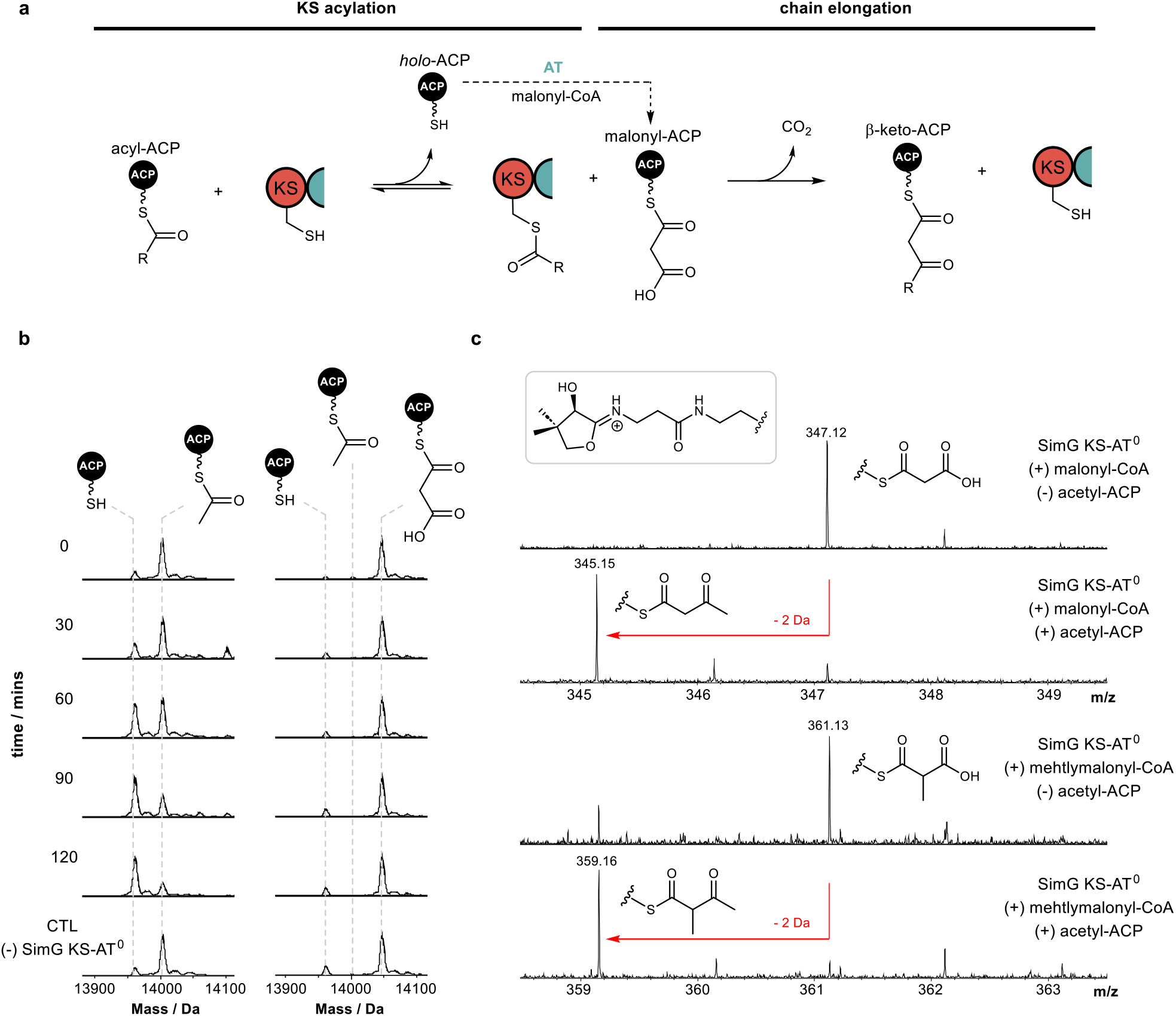
Selectivity of the SimG KS domain towards biosynthetic intermediates. **a**). Schematic of KS-catalysed transacylation and chain elongation steps. **b**). Deconvoluted mass spectra of SimG acetyl-ACP (*left*) and malonyl-ACP (*right*) following incubation with SimG KS-AT^0^ over a 120 min time course. Conversion of the acetyl-ACP species to *holo*-ACP indicates transfer to the KS domain (see **Supplementary Figure. S6** for KS acylation data), whereas negligible transfer is observed for the malonyl-ACP species. **c**). Ppant ejection spectra of chain elongation assays for acetyl-ACP and malonyl-ACP (*top*) and acetyl-ACP and methylmalonyl-ACP (*bottom*). Compared to acetyl-ACP-free controls, Ppant ejection ions corresponding to chain elongated products were detected for acetoacetyl-ACP (*m/z* = 345.1) and 2-methylacetoacetyl-ACP (*m/z* = 359.1). All experiments were conducted in triplicate and representative spectra are shown.

Using our aforementioned KS-AT^0^ construct, whereby the AT domain has been inactivated to prevent competing reactions, we first examined the ability of the KS domain to be transacylated with the acetyl starter unit from the ACP domain, required to prime the biosynthesis of Hma (**3**). Here, incubation of a 1:1 ratio of acetyl-ACP to KS-AT^0^ didomain resulted in complete loss of the acetyl unit from the ACP domain over a 2 hr time period (**Fig. 5b**). Despite complete conversion of the ACP domain to the *holo* form, only ∼35 % acylation of the KS-AT^0^ was observed at the reaction end point (**Supplementary Fig. S6**), suggesting that the acetyl unit is slowly hydrolysed from the KS domain if not utilised for a chain extension reaction. To check whether acetyl-CoA can directly prime the KS domain, rather than via the AT domain, an excess of acetyl-CoA was incubated with the KS-AT^0^ didomain, however no acylation was observed (**Supplementary Fig. S6**), indicating that the AT domain is solely responsible for priming the hrPKS. Incubating the KS-AT^0^ didomain with the extender unit malonyl-ACP species resulted in no loss of the malonyl unit over the time course of the reaction, and importantly, no decarboxylation to the acetyl-ACP species was observed which could serve as a starter unit (**Fig. 5b**, *right*). However, extended incubations of malonyl-ACP in the absence of KS-AT^0^ resulted in low levels of acetyl-ACP formation (**Supplementary Fig. S7**), which accounts for previous observations of *in vitro* product formation when providing hrPKSs with solely malonyl-CoA.^28,31,39–43^

Having established that the KS domain can be acylated by the acetyl-ACP species, we next examined its ability to catalyse a chain extension reaction. Due to the small mass difference between the malonyl-ACP substrate and the acetoacetyl-ACP product (ΔM = −2 Da), we elected to monitor the reaction using Ppant ejection, allowing an unambiguous mass shift associated with chain elongation to be observed.^44,45^ Here, incubation of SimG KS-AT^0^ with a mixture of acetyl- and malonyl-ACP resulted in near-complete consumption of the malonyl-ACP (*m/z* = 347.16) and emergence of a new peak corresponding to the elongated acetoacetyl-ACP product (*m/z* = 345.15) (**Fig. 5c**, *top*). Omission of acetyl-ACP or the KS-AT^0^ didomain from the reaction resulted in no product formation, highlighting that both the acyl-donor and KS domain are required for a productive elongation reaction (**Fig. 5c**, *top*). Replacement of malonyl-ACP with methylmalonyl-ACP also led to a productive chain extension reaction, indicated by conversion of the methylmalonyl species (*m/z* = 361.14) to a 2-methylacetoacetyl species (*m/z* = 359.16) (**Fig. 5c**, *bottom*). This indicates a degree of flexibility in the KS domain towards the extender unit, which could be exploited to produce polyketide derivatives. However, two factors likely prevent the KS domain from being presented with a non-native extender unit: i). the gatekeeping activity of the AT domain, which prevents transfer of methylmalonyl units to the ACP domain (**Fig. 3e**), and ii). that (2S-methylmalonyl-CoA is not known to be produced by fungi.^46^

## DISCUSSION

Whilst a large proportion of fungal hrPKSs load acetyl- and malonyl-CoA as their respective starter and extender units, the molecular basis for how they are processed has remained unresolved. Our *in vitro* reconstitution experiments, combined with intact protein MS, have permitted direct monitoring of all steps during the priming and first chain extension reactions for a fungal hrPKS, allowing proposal of a viable catalytic route likely applicable to all hrPKSs (**Fig. 6**).

**Figure 6.**
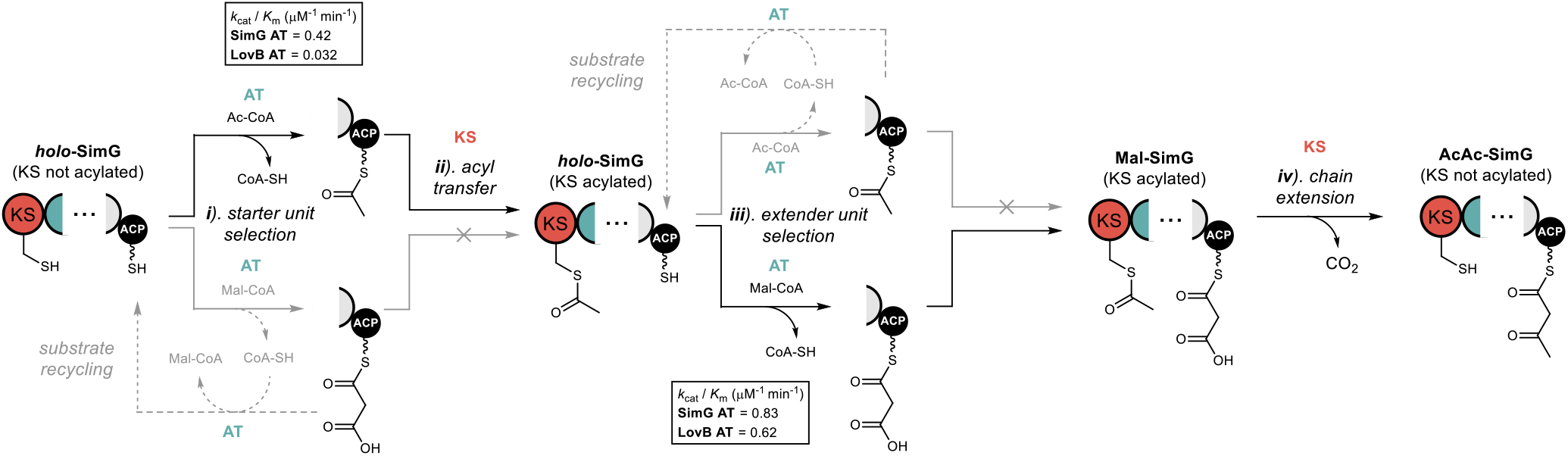
Biosynthetic logic of acyl group selection in fungal hrPKSs. Schematic depicting possible routes for acyl groups during priming of a fungal hrPKS. The forward course of the reaction is shown in black arrows, and substrate recycling events with grey arrows. Defined steps are numbered i – iv, and are referenced in the main text. The *k*_cat_ / *K*_m_ values for acetyl and malonyl loading events by the LovB and SimG AT domains are shown.

Starting from a *holo*-hrPKS state, the AT domain must select an acetyl-CoA starter unit from the cellular pool, competing with malonyl-CoA extender units and other acyl-CoA substrates (**Fig. 6**, step i). The AT domain exhibits a preference for malonyl-CoA over acetyl-CoA (2-fold for SimG, 20-fold for LovB), meaning that extender units are likely erroneously loaded onto the ACP domain in the initial stages. It is worth noting that the intracellular concentration of acetyl-CoA is typically higher than malonyl-CoA, and it is tempting to speculate that the AT domain may have evolved a preference for malonyl-CoA to offset this difference.^47^ Whilst an ‘incorrect’ malonyl unit can be loaded at this point, the KS domain acts as a proofreader: it will not accept a malonyl-ACP as a starter unit, nor can it catalyse decarboxylation at a sensible rate to yield an acetyl-ACP starter unit. Therefore, the AT domain must utilise free CoA to remove erroneous malonyl units from the ACP domain. This process of loading / offloading continues until an acetyl starter unit is selected, which can prime the KS domain. The offloading of acetyl units appears to be less efficient than malonyl units (**Fig. 4d**), this may allow sufficient ‘dwell time’ for transfer to the KS domain to occur (**Fig. 6**, step ii).

The mechanism by which the KS domain discriminates between acetyl- and malonyl-ACP at this step is currently unclear and requires further study. Recent structural work on the mFAS KS domain highlighted that two phenylalanine residues, Phe200 and Phe395, contribute to substrate specificity by sterically blocking under-processed intermediates (i.e. β-keto, β-hydroxy and α,β-unsaturated) from acylating the active site cysteine.^48^ Equivalent positions in the SimG KS domain are Val254 and Ile452, respectively, and an AlphaFold structural model of the SimG KS domain suggests that they do not provide the required steric hinderance to block branching at the β-carbon (**Supplementary Fig. S8**). Further factors such as allosteric regulation within the condensing region^49^, and / or the possibility of substrate control by the ACP domain^50,51^, may also impact this transfer step and require further investigation. Our data has also highlighted that whilst the AT domain can become acylated with a variety of medium / long chain acyl-CoAs present in the cellular milieu (an observation also made for the mFAS AT domain^35,48^), transfer to the ACP domain is only permitted for acetyl and malonyl units (**Fig. 3e**). Although transfer to the ACP domain is blocked, and likely prohibits the use of non-native starter units, offloading back to free CoA prevents the AT domain from being irreversibly inhibited (**Supplementary Fig. S5**). This is analogous to the editing mechanism employed by aminoacyl-tRNA synthetases^52^, and highlights a previously unappreciated proofreading role played by AT domains for ensuring biosynthetic fidelity.

Following acetyl transfer to the KS domain, the *holo*-ACP then needs to be loaded with a malonyl extender unit (**Fig. 6**, step iii). Despite the intrinsic preference of the AT domain for malonyl, another cycle of substrate loading / offloading can occur at this stage, to ensure a malonyl unit, instead of an acetyl unit is loaded onto the ACP domain. Subsequently, the KS domain catalyses a decarboxylative chain extension reaction between the acetyl-KS and malonyl-ACP species, resulting in the acetoacetyl-ACP product (**Fig. 6**, step iv). Interestingly, our data indicates that the KS domain lacks specificity towards the extender unit, allowing chain extension to be catalysed with the non-native methylmalonyl-ACP species. Whilst fungi are not known to produce methylmalonyl-CoA^46^, meaning exposure of the KS domain to this particular extender unit is unlikely, and further underscores the gatekeeping role of the AT domain, which prevents transfer of a methylmalonyl unit onto the ACP domain (**Fig. 3e**). The inherent flexibility of the KS domain towards extender units could be exploited through AT domain swapping experiments, utilising previously characterised alkylmalonyl-CoA accepting AT domains.^53–55^ It might be expected that the KS domain exhibits specificity during the acylation step, only selecting for the correct biosynthetic intermediate from the modifying cycles. Our data shows the first indications of this specificity, with the SimG KS domain unable to be acylated by the incorrect malonyl-ACP species (**Fig. 5b**), and will be the focus of further work on this system.

Taken together, this work unequivocally demonstrates the trajectory of acyl groups during the chain priming and elongation process in fungal hrPKSs. We show that the KS and AT domains of the condensing region are reliant upon each other, working closely together to ensure biosynthetic fidelity of the system. During the priming step, the role of the KS domain is to proofread the cargo appended to the ACP domain, allowing acylation with the starter unit but not the extender unit. However, once acylated, the KS domain is unable to discriminate between extender units for the elongation reaction. Here, the AT domain ensures that the KS domain is always presented with the correct extender unit for chain elongation, with the selectivity of the AT domain confining the system to one starter / extender unit pairing.

Using this biosynthetic framework, it seems likely that hrPKSs which incorporate non-acetyl starter units must also use the AT domain to select the correct acyl units. For example, the hrPKS-NRPS responsible for pseurotin A production, PsoA, has been shown to preferentially use propionyl-CoA as the starter unit over acetyl and other acyl-CoAs.^31^ Here, our data would imply that the PsoA AT domain is responsible for loading both the propionyl and malonyl units required for chain assembly. This can also be extended to AT domains that use benzoyl-CoA as the starter unit (e.g. in strobilurin and squalestatin S1 hrPKSs), which appear to be phylogenetically discrete from other acetyl-incorporating AT domains.^33^ The molecular basis for how some AT domains are able to select for non-acetyl starter units, whilst maintaining their ability to recognise malonyl extender units, remains to be elucidated and will be the focus of future studies. Overall, this work provides a basis for future research to decode the intrinsic programming of these enigmatic enzymes and lays the foundations for future bioengineering approaches aimed at creating novel chemical scaffolds.

## ACKNOWLEDGEMENTS

This work was supported by a UKRI Future Leaders Fellowship to M.J. (MR/W011247/1), from which N.B.A. and Y.T.C.H were funded. M.E.F gratefully acknowledges funding from an EPSRC Doctoral Training Partnership (EP/T51794X/1) studentship. Y.H. and S.L. were supported by an NIH grant (R35GM151205). The Bruker MaXis II instrument used in this study was funded by the BBSRC (BB/M017982/1). The authors are grateful to Professor Yi Tang for use of the pXW55-SimG plasmid, and for valuable discussions related to the work.

## AUTHOR CONTRIBUTIONS

M.J. and M.E.F. conceived and designed the study. M.J., M.E.F. and Y.H. generated *E. coli* and yeast expression constructs. N.B.A, S.L. and Y.H. performed heterologous metabolite production. M.E.F., Y.T.C.H. and M.J. conducted protein overexpression and purifications. M.E.F. constructed mutant / deletion plasmids, carried out all biochemical assays on recombinant proteins, and constructed AlphaFold models. M.J. wrote the manuscript with input from all authors.

## DATA AVAILABILITY

The minimum dataset required to interpret, verify and extend the work is provided in the manuscript and supplementary information. The raw data for **Figs. 3, 4** and **5**, and **Supplementary Figs. 1, 2**, and **4 – 7**, which were processed via standard deconvolution, are available upon written request to the corresponding author. The DNA sequence of the full-length SimG construct, and associated protein sequences of excised domains used in this study, are reported in the supplementary information. A reporting summary for this Article is available as a supplementary information file.

