## Supplementary Information for "Biochemical dissection of a fungal highly reducing polyketide synthase condensing region reveals basis for acyl group selection"

### 1. Supplementary Figures

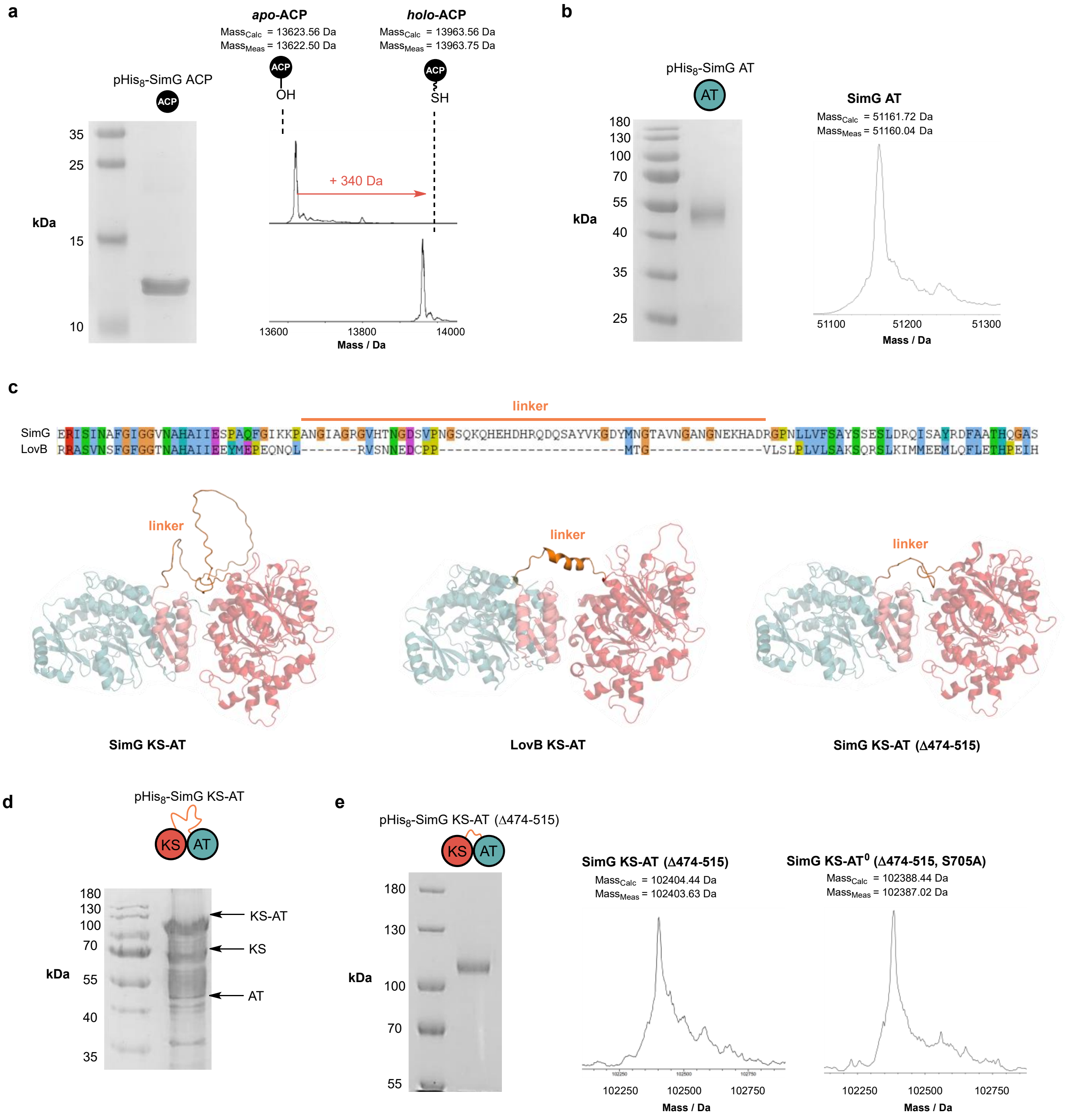


**Supplementary Figure S1.** Biochemical dissection of the SimG hrPKS condensing region. **a**). 12 % SDS-PAGE gel of purified SimG ACP domain (*left*) and deconvoluted mass spectra of *apo*- and *holo*-ACP domain (*right*). **b**). 10 % SDS-PAGE gel of purified SimG AT domain (*left*) and deconvoluted mass spectrum (*right*). **c**). Sequence alignment of the linker region between KS and AT domains in SimG and LovB hrPKSs (*top*) and structural visualisation of the linker regions (*bottom*). **d**). 8 % SDS-PAGE gel of SimG KS-AT didomain showing degradation after $Ni^{2+}$-NTA purification. **e**). 6 % SDS-PAGE gel of purified SimG KS-AT(Δ474-515) (*left*), deconvoluted mass spectra of KS-AT (Δ474-515) (*middle*) and KS-AT (Δ474-515, S705A) (*right*).

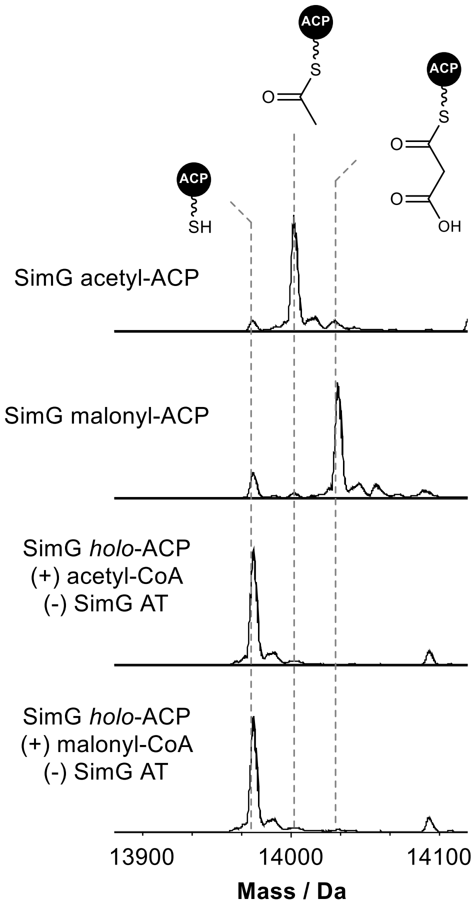


**Supplementary Figure S2.** Deconvoluted mass spectra of SimG ACP species from control reactions of SimG AT-catalysed transfer of acetyl and malonyl units. No transfer of acetyl or malonyl units was observed over the duration of the reactions when the SimG AT domain was omitted.

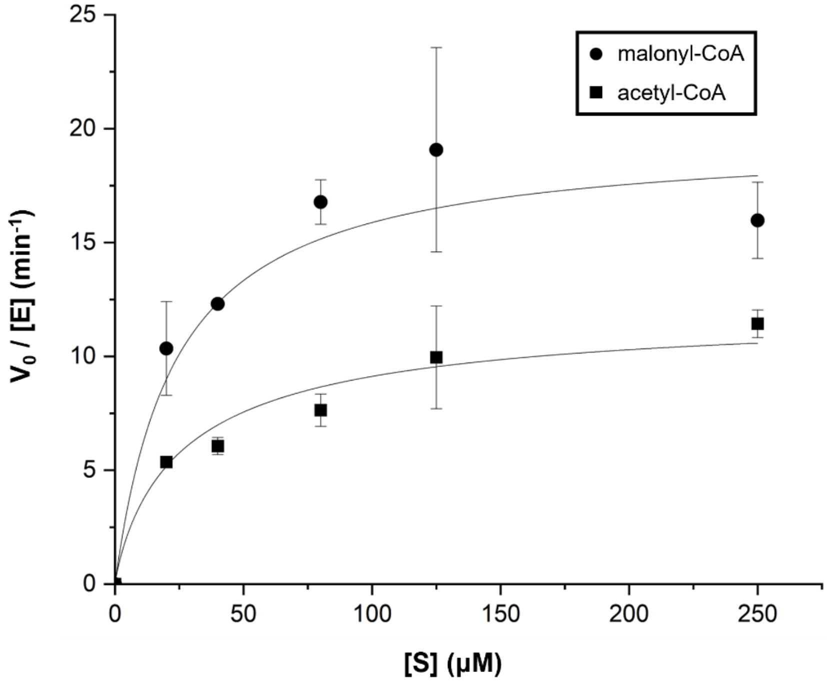


**Supplementary Figure S3.** Michaelis–Menten plot of SimG AT-catalysed transacylation of acetyl- and malonyl-CoA to SimG *holo*-ACP domain. Assay conditions used 1 µM SimG AT domain and 50 µM SimG holo-ACP domain, with acyl-CoA concentrations between 20 – 250 µM. The kinetic constants are shown in the main text (**Fig. 3d**).

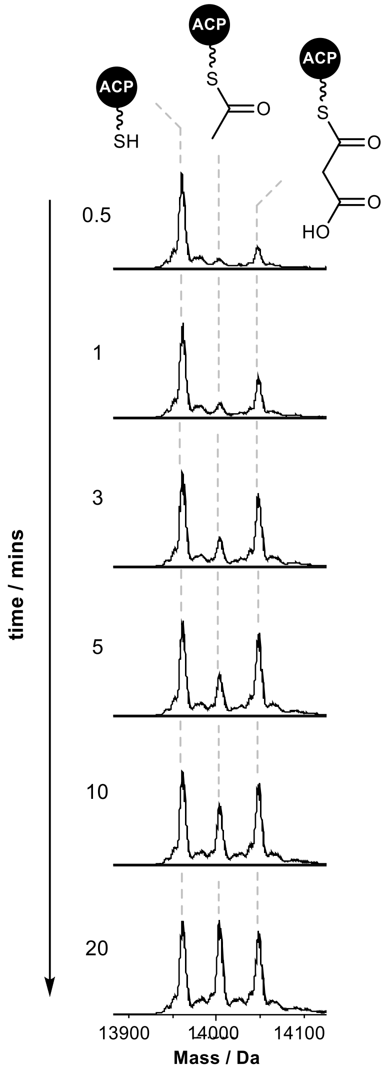


**Supplementary Figure S4.** Deconvoluted mass spectra of SimG ACP species from a time-course analysis of SimG AT-catalysed loading of acetyl- and malonyl- units in a competition reaction. Early time points show preferential loading of malonyl, before accumulation of acetyl units becomes the major species. Faster offloading of malonyl units compared to acetyl units (see **Fig. 4**) allows the acetyl-ACP species to persist over time.

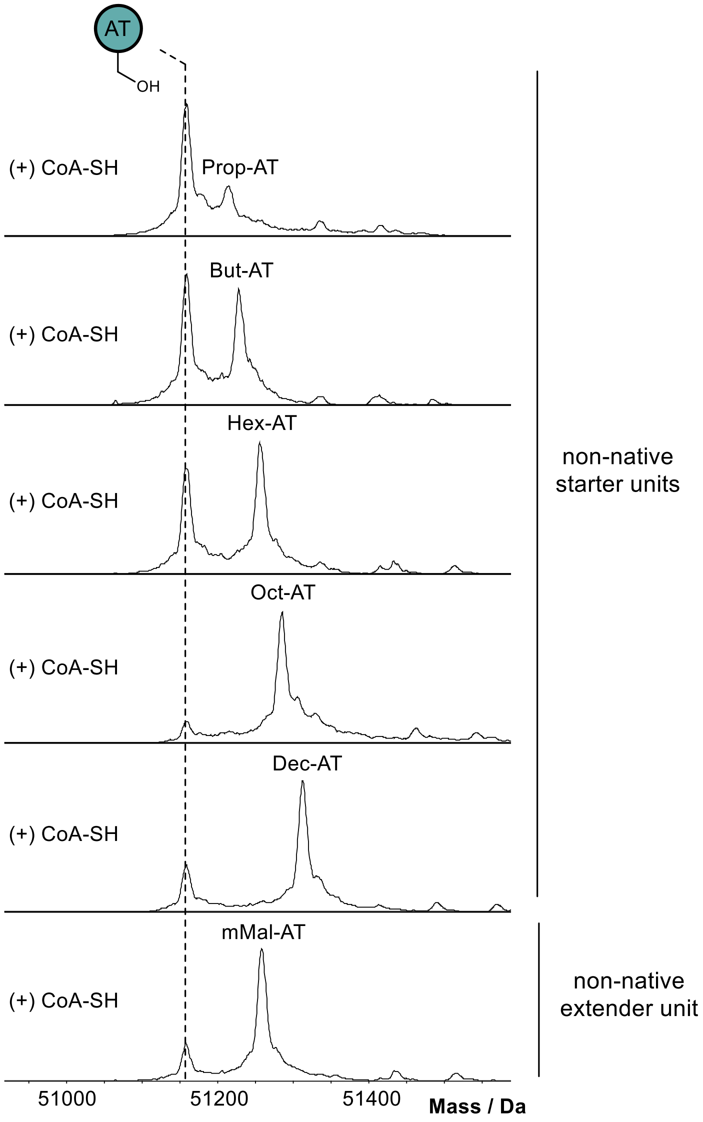


**Supplementary Figure S5.** Deconvoluted mass spectra of non-native acyl-SimG AT species following offloading reactions. A decrease in offloading efficiency is observed as chain length increases for non-native starter units (propionyl-, butyryl-, hexanoyl-, octanoyl-, decanoyl-AT), whilst negligible offloading was observed for the non-native extender unit, methylmalonyl(mMal)-AT.

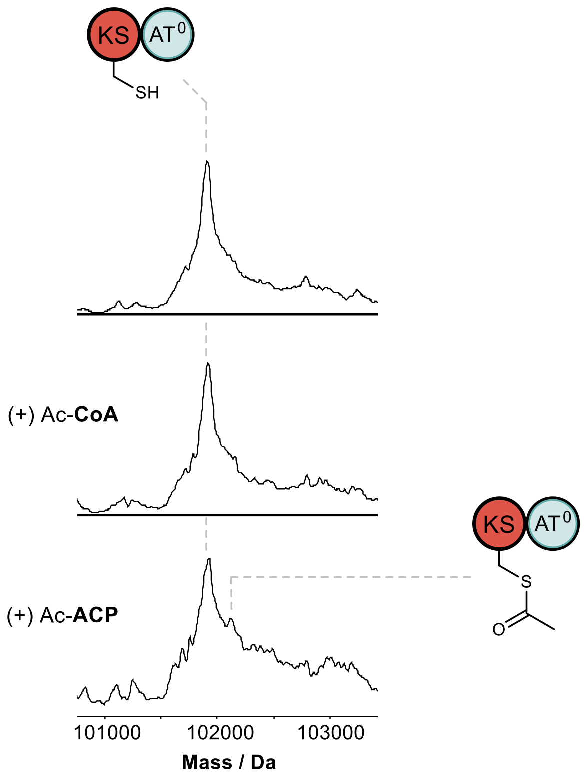


**Supplementary Figure S6.** Deconvoluted mass spectra of SimG KS-AT$^0$ (*top*), following incubation with excess acetyl-CoA (*middle*) and acetyl-ACP (*bottom*). No acetyl transfer to the KS domain is observed using acetyl-CoA, whereas a peak corresponding to +42 Da can be observed when acetyl-ACP is supplied. Full acetylation of the KS domain is likely prevented by competing hydrolysis during the course of the reaction.

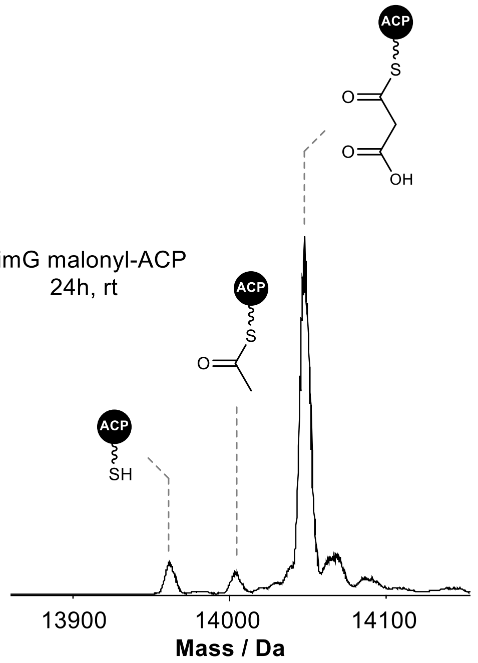


**Supplementary Figure S7.** Deconvoluted mass spectrum of malonyl-SimG ACP following a 24 h incubation in storage buffer (see Methods) at room temperature. Spontaneous decarboxylation of the malonyl-ACP species gives rise to a small amount of acetyl-ACP, which can serve as a starter unit.

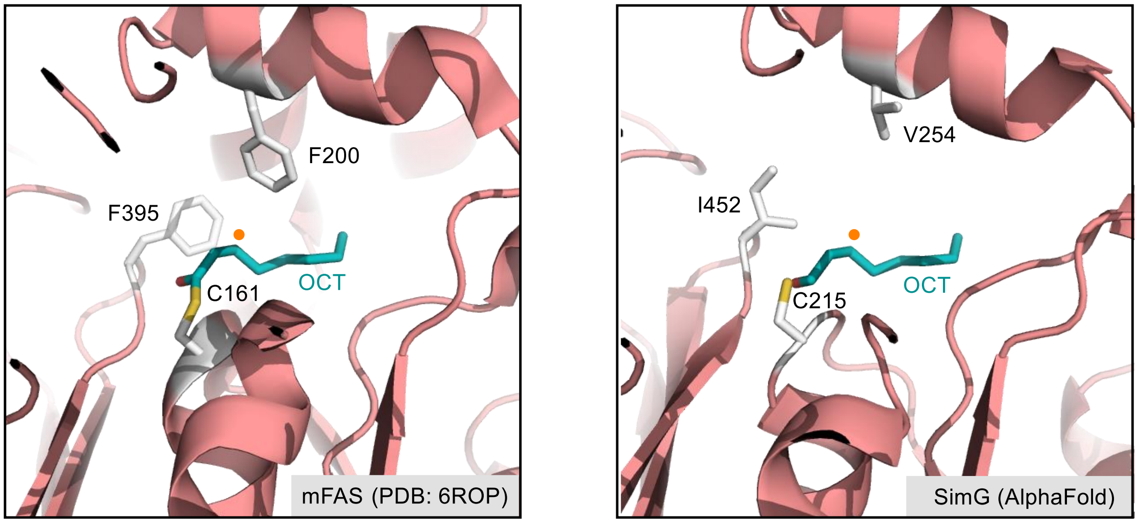


**Supplementary Figure S8.** Structural comparison of KS domains from mFAS (*left*) and SimG (*right*). The mFAS structure (PDB: 6ROP)[1] has an octanoyl group covalently bound to the active site C161 residue, with branching at the β-position (indicated by an orange circle), or a conformationally restricted α,β-unstatured intermediate, prohibited via steric hinderance from F200 and F395. These residues are replaced with V254 and I452 in an AlphaFold model of the SimG KS domain, indicating increased space within the active site.

### 2. Methods

#### 2.1. Molecular Cloning and Mutagenesis

*2.1.1. SimG hrPKS*

The SimG yeast expression plasmid pXW55-SimG was constructed using *in vivo* yeast recombination cloning. The intron-less SimG gene was cloned from a cDNA library prepared from *Tolypocladium inflatum* NRRL 8044. Briefly, mRNA was extracted from *T. inflatum* mycelium by using the PureLink™ RNA Mini Kit (Invitrogen) following the manufacturer's instructions. The cDNA library was then syntheszied by using the SuperScript III FirstStrand Synthesis System (Invitrogen) with Oligo-dT primers. The SimG gene fragments were then amplified and assembled with the linearized XW55 vector in *S. cerevisiae* JHY686 strain.[2] The resulting plasmid was recovered by Zymoprep Yeast plasmid miniprep for propagation in *E. coli* XL-10 cells and verified by sequencing.

*2.1.2. SimG KS-AT, AT and ACP*

The amplifications of SimG KS-AT didomain, SimG AT domain, and SimG ACP were performed from the full SimG hrPKS construct (2.1.1.) using Q5 DNA polymerase (NEB) and the primers detailed in **Table S2**. PCR products were confirmed on a 1 % agarose gel prior to digestion using NdeI and HindIII restriction enzymes (NEB) and ligation into modified pET28a(+) vector using T4 ligase (NEB) which had also been digested with NdeI and HindIII restriction enzymes (following manufacturers protocols). The modified pET28a(+) vector included two additional His residues in the His-tag and a G2K point mutation to reduce gluconylation of the final protein product (see corresponding protein sequence in **Section 3**). The ligation mixture was used to transform *E. coli* TOP10 cells (Invitrogen) which were plated onto LB agar containing kanamycin (50 μg/mL) and grown at 37 ˚C overnight. Colonies were selected and grown overnight in LB media containing kanamycin (50 μg/mL) prior to plasmid isolation using a miniprep kit (Thermo). The integrity of the gene inserts was confirmed by Sanger Sequencing (Eurofins).

*2.1.3. SimG KS-AT (Δ473-514)*

The amplification of SimG KS-AT didomain (Δ473-514) was performed from the SimG KS-AT construct (2.1.2.) using Q5 DNA polymerase and the primers detailed in Table S2. PCR product was confirmed by 1 % agarose gel prior to ligation using a KLD reaction kit (NEB) following manufacturers protocols. The ligation mixture was used to transform *E. coli* DH5α cells (NEB) which were plated onto LB agar containing kanamycin (50 μg/mL) and grown at 37˚C overnight. Colonies were selected and grown overnight in LB media containing kanamycin (50 μg/mL) prior to plasmid isolation using a miniprep kit (Thermo). The desired deletion was confirmed by Sanger Sequencing (Eurofins).

*2.1.4. SimG KS-AT$^0$ (Δ473-514), (S705A)*

SimG AT domain containing the S705A point mutation (AT$^0$) was gene synthesised in the pBSK vector (Epoch Life Sciences) for use in this work. The amplification of the SimG AT$^0$ fragment was performed from the gene synthesised construct using Q5 DNA polymerase (NEB) and the primers detailed in Table S2. The amplification of the pET28a(+) SimG KS fragment was performed from the SimG KS-AT (Δ473-514) construct (see Section 2.1.3) using Q5 DNA polymerase (NEB) and the primers detailed in **Table S2**. PCR products were separated on a 1 % agarose gel and excised bands were purified using a gel extraction kit (Thermo). The purified PCR products were ligated using a NEBuilder® HiFi DNA Assembly kit (NEB) following manufacturers protocols. The ligation mixture was used to transform *E. coli* DH5α cells (NEB) which were plated onto LB agar containing kanamycin (50 μg/mL) and grown at 37˚C overnight. Colonies were selected and grown overnight in LB media containing kanamycin (50 μg/mL) prior to plasmid isolation using a miniprep kit (Thermo). The presence of the point mutation was confirmed by Sanger Sequencing (Eurofins).

#### 2.2. Production of 3*R*-hydroxyl-4*R*-methyl-6*E*-octenoic acid in *S. cerevisiae* JHY686

For the production of 3R-hydroxyl-4R-methyl-6E-octenoic acid, *S. cerevisiae* JHY686 strain[2] harboring the pXW55-SimG plasmid was inoculated to 4 mL of Yeast Synthetic Drop-Out medium without uracil. The cells were grown for 72 hours with constant shaking at 28 ºC. A 1 mL aliquot of the seed culture was inoculated to 1 L YPD medium supplemented with 1% dextrose. The culture was shaken at 28 ºC for 72 hours. The cells were removed by centrifugation and the liquid medium was acidified with formic acid, followed by extraction with ethyl acetate. The organic layer was concentrated *in vacuo* and the title compound was purified by Biotage Isolera chromatographic system.

#### 2.3. Protein overexpression and purification

An aliquot of chemically competent *E. coli* BL21 Star (DE3) cells (50 µL) was transformed with relevant plasmid DNA. Transformed cells were plated onto LB agar plates containing kanamycin (50 µg/mL) and incubated overnight at 37˚C. A single colony was picked and used to inoculate LB media (10 mL) containing kanamycin (50 µg/mL) and incubated overnight at 37˚C with shaking (180 rpm). The overnight culture was used to inoculate a flask of LB media (1 L) with kanamycin (50 µg/mL) which was left to grow at 37˚C with shaking (180 rpm) until an $OD_{600}$ of 0.7 - 1.0 was reached. Protein expression was induced by the addition of IPTG (250 µM) and incubated at 15˚C with shaking (180 rpm) overnight.

Cells were centrifuged (4000 rpm, 15 minutes, 4 ˚C) and resuspended in loading buffer (20 mM Tris HCl, 100 mM NaCl, 20 mM imidazole, pH 8.0). The cells were lysed by high pressure (20 psi) cell disruption and centrifuged (17,000 rpm, 30 minutes, 4˚C) to pellet the insoluble cell components. The cell lysate was filtered through a 0.45 µm syringe filter and loaded onto a 1 mL HiTrap™ Nickel FastFlow column. Following a wash with loading buffer (10 mL), the protein was eluted from the column with buffers of increasing imidazole concentrations (5 mL of 50 mM, 3 mL of 100 mM, 3 mL of 200 mM, 3 mL of 300 mM). Fractions were checked for the protein of interest by SDS-PAGE and those containing protein were concentrated and exchanged into storage buffer (20 mM Tris HCl, 100 mM NaCl, pH 7.6) using an appropriate size Vivaspin centrifugal concentrator (4000 rpm, 4 ˚C). Once concentrated to approx. 1 mL, 50 µL aliquots of protein were flash frozen in liquid $N_2$ and stored at -80 ˚C.

#### 2.4. Conversion of *apo*-SimG ACP to *holo*- / acyl-SimG ACP

SimG ACP (200 µM) was converted to its *holo*- or *acyl*- form by incubation in storage buffer (20 mM Tris base, 100 mM NaCl, pH 7.6) with 10 mM $MgCl_2$, 2 µM Sfp PPTase, and 1 mM CoA/*acyl*-CoA in a total of 50 µL for 30 minutes at 25 ˚C. For use in **Sections 2.7 – 2.9**, the generated ACP species as diluted 5-fold with storage buffer and concentrated to approx. 50 µL using a 10 kDa MWCO Vivaspin centrifugal concentrator (12000 rpm, 4 ˚C) to reduce the amount of CoA present in the subsequent assay.

#### 2.5. SimG AT acylation and ACP transfer

To investigate AT acylation, SimG AT domain (50 µM) was incubated in storage buffer (20 mM Tris base, 100 mM NaCl, pH 7.6) with 1 mM *acyl*-CoA in 50 µL at 25 ˚C. The reaction was quenched with 1 % formic acid at 1 minute following AT domain incubation. Samples were diluted 5-fold with milliQ $H_2O$ and analysed by UHPLC-ESI-Q-TOF-MS. To capture ACP transfer, SimG *holo*-ACP (50 µM) was incubated in storage buffer (20 mM Tris HCl, 100 mM NaCl, pH 7.6) with 1 mM acyl-CoA and 10 µM AT domain in a total of 50 µL at 25 ˚C. The reaction was quenched with 1 % formic acid at 1 minute following AT domain incubation. Samples were diluted 5-fold with milliQ $H_2O$ and analysed by UHPLC-ESI-Q-TOF-MS.

#### 2.6. SimG AT substrate offloading

SimG acetyl-ACP or malonyl-ACP (50 µM) was incubated in storage buffer (20 mM Tris base, 100 mM NaCl, pH 7.6) with 1 mM CoA and 10 µM AT domain in a total of 50 µL at 25 ˚C. Reactions were quenched with 1 % formic acid after 1 minute following AT domain incubation. Samples were diluted 5-fold with milliQ $H_2O$ and analysed by UHPLC-ESI-Q-TOF-MS.

#### 2.7. SimG acyl-ACP chain transfer

SimG acetyl / malonyl-ACP (50 µM) was incubated in storage buffer (20 mM Tris HCl, 100 mM NaCl, pH 7.6) with KS-$AT^0$ (50 µM) in a total of 100 µL at 25 ˚C. For *ace*-ACP and *mal*-ACP, the reaction was quenched with 1 %

formic acid at 0, 0.5, 1, 1.5, 2, 2.5, and 3 hours following KS-$AT^0$ incubation. Samples were diluted 5-fold with milliQ $H_2O$ and analysed by UHPLC-ESI-Q-TOF-MS.

#### 2.8. SimG acyl-ACP chain extension

SimG malonyl-ACP (50 μM) and acetyl-ACP (50 μM) were incubated in storage buffer (20 mM Tris HCl, 100 mM NaCl, pH 7.6) with KS-$AT^0$ (100 μM) in a total of 50 μL for 30 minutes at 25 ˚C. Samples were diluted 10-fold in milliQ $H_2O$ and analysed by UHPLC-ESI-Q-TOF-MS.

#### 2.9. SimG AT acyl-CoA competition

SimG *holo*-ACP (50 μM) was incubated in storage buffer (20 mM Tris HCl, 100 mM NaCl, pH 7.6) with 125 μM acetyl-CoA, 125 μM malonyl-CoA and 1 μM AT domain in a total of 100 μL at 25 ˚C. The reaction was quenched by addition of 1 % formic acid at 0.5, 1, 3, 5, 10, and 20 minutes following AT domain incubation. Samples were diluted 5-fold with milliQ $H_2O$ and analysed by UHPLC-ESI-Q-TOF-MS.

### 3. Sequences, Tables and Accession Numbers

#### 3.1 DNA Sequences

**>simG_hrPKS (minus START codon, as cloned into XW55 vector)**

```
GCTCCTCAACAAATATCACCCCCCAATGGGACGGCATCCCCAGAGGCCGGGACTCCAAGATATACGTACAGACCAGCTTCACCTGCACCAGAATCGCCATGGGCAGACAGCGCGGAAG
GGGCCGAGTTGGCCAAGCCCATGGCCATCATCGGCATGGCCATGCGTCTCCCGGGAAGCGTACGCTCGGGTGACGACCTCTGGGACCTCCTCTCCACGAGGAAGAGCGGACTCTGCGA
TATCCCCAAGAACCGGTTCAACGCCGACGGGTTCTACGACCCGGCTCGGGGCCCCGGGACGATCCCGGTCAAGAAGGGATATTTCCTCCATGACGTGCAGATCGAGGAGTTCGACACC
AACGTCTTCCCCATCCCCAAGATGGAGCTGGAGCGCCTGGATCCTGCCCAGCGACAGTTGCTCCAGGTAGCCTACGAGTGCATAGAGAACGCAGGCGGGACCTCGTGGCGTGGGAGCA
GAACTGGGTGCTACATCGGCGAGTTCGGCGAGGACTTTGCCGACAGTTCTGCACGCGAGTCACAGCAAAGGGGCAACTTACGATACACCGGCCTTGCCGATTTCGCAATAGCCAACAG
GATCTCGTATGAGCTCGACCTTCAAGGCCCGAGTATGGTGGTCAAGACGGCTTGCTCGTCTTCCCTCGTGTGCCTCGACCTGGCCTGCAAGGCCATCCAGAGCGGCGAATGCGAAAGT
GCCCTCGTCGGCGGCGTCAGTATGCTCTTCTCGCCCGCCACGTACATCAGCGTGACCGACCTGGGCGTCATCTCGCCCAACGGCCAGTGTCGGAGCTTCGACGCCGGCGCGGACGGCT
ACGCGCGAGGGGAGGCCGTCAACATGGTCATGATCAAGAGGCTAGATCACGCTCTGAGGGACAAGGACCCCATCCGTGCCGTCATCCGCGGGACCGGTGTCAACACCGACGGCAGAAC
CAACGGCATGCTGACCCCAAGCTCCGCCGTGCAAGCGGATCTCATCCGGCACACATACGAGATCGCTGGAATAGATGACCTGTCACAAACCGCCGTCGTGGAGTGTCACGGCACCGGA
ACTCCGGTCGGCGACCCCATCGAGACGGAGGCTGTCGGACGGTGCTTTGGCGACGACGGCGTTACCATCACATCAGTTAAACCAAATCTTGGACACGCTGAAGCTGCCGCCGGCCTGT
CCAGCCTGATCAAGTGTGTCCTGAGCTTGGAGCACCGGCAAGTGCTGCCAAATATCAACTTCGAGACACCGAATCCAGACATTCCGTTTGAGAAATACAAACTCCGCGTCCCGACCGA
GGTTGAATCTTGGCCGCAGGGTCGAGCAGAGAGGATAAGCATCAACGCCTTCGGCATCGGAGGCGTCAACGCGCACGCAATCATCGAATCCCCGGCGCAGTTTGGAATCAAGAAGCCT
GCCAACGGCATCGCCGGCCGCGGCGTACACACGAACGGCGATTCTGTTCCCAATGGTTCTCAAAAGCAGCATGAGCATGATCACCGTCAAGACCAAAGCGCCTACGTGAAGGGCGATT
ATATGAACGGCACTGCCGTCAACGGGGCAAACGGCAACGAAAAGCATGCTGACCGCGGACCGAATCTCCTGGTCTTCTCCGCCTATAGCTCCGAGTCTCTCGATCGGCAGATCAGTGC
TTACCGTGACTTTGCAGCAACTCACCAAGGCGCCTCGCTCAAGGATCTGGCATATACCTTGTCGAGTCGGCGCGATCACAGGCCCTACCGGGCTTATGCCATCGCCGACGACGCATCC
GGTGTCCAAGACGCCTCCGCCACCGTTAACGCCGTCGTAGACGGAGCACCTCCGCCGGTGGGCTGGATCTTCACCGGACAGGGTGCGCAGTGGCCGGAGATGGGCGCCAGGCTGATCG
ATACCAGTGCTGCGTTCAGAAACAGGATCCGTAAGCTGGACAAGTACCTGCAGACGCTGAAGCTGGAGCCACCGGTGAGCATTGAAGCCGAGCTGCGCAAGACCAAGGAAGACAGTCG
AGTGCACCGTCCCGAGATGGGACATCTCGTATGCGTCGCCATGCAGGTAGCCCTGGTCGACGTCCTCCGCTCCTGGAACATCGTCCCAGACTTTGTCCTGGGCCACTCGTCGGGCGAA
ACAGCCGCCGCCTACGCCTGCGGAGCCATCACGGCGGAGGCTGCCGTATACGCGGCAACGAGACGAGGCATCGGCAACGCATCTTCCCAGAGGAAAGGCTCCATGGCTGCCGTCGGGC
TGGGCAGGGATGAGATACAGCCGTTCCTGCTCGAGGGGGTCAACATCGCCTGCGAGAACAGTCAGAGCAACGTCACGCTGTCGGGAGATACGGAGCAGGTTGAGAGCATCGTAGCGAC
GCTCAAGGCCGAGAGACCGGGGGTCTTTGCTCGTTTGCTCCGGGTAGAGAAGGCGTATCACTCCCATCACATGTGTGAATACGGGCCGGTGTATGAGGAGCAGCTGAAGCCCGTGGTT
CGCAGCGCTGACCCCGAGATCCCATTCTACTCCTCCGTCACCGGTGACCGCCTGACCGGCGAGGGACAGCTCGGGGCATCGTATTGGCGGGCGAACATGGAGAACACGGTTCTCTTCA
ACCCGGCGCTCCGCTCCGCCCTCCGAGATCGACCCGGAAAGCTGGTCCTAATCGAGTTGGGACCTCATCCCGCCCTTGAAGGACCAGTCGGCCAGATTCTCCGAGACCTGGGTCGCAC
AGACGACATCTACCTTGGGACCTGTATCAGGGGCAGCGAGTGCGATCGCAGCCTGCTTCACCTCGGCGGAAAGCTGTACCAGCAGTCGATACCGATGGACCTTGCAGCGATCTGCCCG
CCCGGCGCCGTCTTGACGAACCTGCCGCGCTATGCCTGGAGCCAGGACACGACGCACTGGGAAGAATCCAGGGTTTCGCGCGGCTGGAGGCTCCGGGAACATCCACCCCACGAGCTTC
TCGGGAGCAGAGTGCTGGAGACGGAGGGCGAGCCGTGTTGGAGAAAGGTAATGGCCCTCGAGGACGCTCTCTGGCTGGATGGCCACGAGGTCAACGGCCAGGTCCTGTTCCCGGCTGC
TGGATACATCTCCATGGTAGGCGAGGCGCTCCGACAGCTGACCGGCGACGCGACATTCACCGTCAGGAACGTCCGCATCATGTCGGGCCTGGTGCTGAGCTCGGACAAGCCGGTCGAG
CTGGTCACTCTATTGAGGTCGACCACGATGGGCACCTCGGACGATTCGGAATGGTACGAGTTCACCATCACCTCGTTCGACGGCACGGCATGGGTCCGGAACTGCCACGGCGAGGCCA
TGGCGTCAAGCGACAAGTCGTTTCACCTCGACTCTGTCTCTCCTGCAGCAGGATCTTTCCCTCGCAAGCTTGACCGGAGGGACTCTTACGACATCCTGAGACGAGTTGGCTTCAACTA
CACCGGGGAGTTTGAAGGGATGGCGGACATCTCGGTCTCGCCGTCGTCGCTTCAAGCCCACGGCGTCACTTCCCCGGCTCGCCAAGTGACCGACAGGCCGTCCAGGTGCGCTACGTAC
TCGGTGCACCCGGCGGTGATCGATCAATGCTTCCAGCTCTTCACGGTGGCCCTATGCCGCGGACTCAGGCGCAGCAACAGGCGGCTGGCGGTCCCGACCTTCATCGAGGAGCTGGTCG
TGTCGCCGAGCGCATCGCCCCTGAGCGTCACCGCCAAGATCAACCGCCTGGACGAGGTGGGATCCTGGACAGGCGATTTCGTTGCCCTCGCGGCCGAGGGGCCCGTCGTGAGGCTCAA
GGGGATGAAGTCCGTCGTCCTCGCGACCCCGGCGCAGTCAGAGCCGCCCCTATCCACGGAACTCGAGTGGAAGCCGCACAGCGACTTTGTCGGATTGGGGGTCGGCCTGCACCCGCGT
GTGCCAAGGAAAAGGGAGTGGGCTCTGCTGGAGGAGCTCGTCCTGCTGTGCAATCTAGACCATCTCGGCCAGATCAAGACCGATGACAGCACTGCGCCGTATCTCATCAAGCTCCTGG
ACTGGATGCGCCTTGTTACGGATCGATACAGGTCCGGCAAGAACCTGTTCGTGTCTGCAGGCGCTGGCCTCGAACATCTGGATCACGACGAGCGAGTCGCCAGGATCGATTTCCTCAT
GGCCCAACTTGCTGACACGCAGGACGTTGTCTTCGCCAGGGCGATCCATCGCCTCTTCATCGAGGCGCCCTCCATATTCGCGGGAGAGGGCCATCCCTTGCACATCCTCATGCCCGAC
AATGTTCTGAGCGACTTTTACGACGCCACATCCTTCGACATGGCCGATGCCGTCCGCCTGGTGGCCAACACGAACCCTCACATGCGCGTGCTCGAGGTTGGCGCGGGAACTGGCAGCA
TGACGGCCAGGCTTCTCCGCGCCTTGACGTCCTCCCACGGCGAGCGCCTCTACAGCCAGTACGGGTACACTGATGTGTCGGCCGGCTTCATGGCCGTGGGGAAGCAGCGCTTTGCCGC
CGTCGAGAACATCGATTACGATGTGCTGGACATATCGAAGGATCCCAAGGAGCAGGGATTCAGGCCCGGCAGCTATGATCTCGTTATCGCTTCCAACGTTCTGCACGCTACCCCTACT
CTGGATGAGACACTTCGCAACGTCTATAGTCTATTGAAGCCCAGTGGACGCTTGTTCCTGGAGGAACTCACTCCAGATGCAATGTTCGTAAACTACGTCATGGGCTTCTTCTACGGCT
GGTGGCTCGGTGCCGAGGACAATCGCGTGGAGCAGCCATGGATCGCGCCCGAATGCTGGGCTGAGAAGTTGGTCGCCGCGGGATTCCAGAAGCCCGAAGCCATCGTTCTGGACAGCGA
CAAGCCATATCAGATCAGCGCGGGGATCATCGCTTCACGGGGGGTACGATCGACCTTGCCCTCCAAGGTGGCGGTTCTGTGCCATACCGCCGATGAGCCCCATGCGGTAGAGATGATC
AGCCGTCTCAGTGCCATGGGCATAGACGTCGAGACATGCCTCTGGGGCCAGCCTCTCCCCTCTTGCGACATCATTTCCGTCCTCGAGCTCGAGAAGCCTTTATTGCATGAAATGTCCG
AGGAGACCTTCAAGACCATCATTGCCTACTTCCAGAGCCACAAGGCAAGGGTGATCTGGGTGACCGAAGCATGCCAGATAGACTGCGCGAACCCCGAGCCTGCAATGATGCTCGGCCT
CGCCAGGACGGTCCGGAACGAATACTCACTACAGATGTACACCGTCGAGCTCGACAAAGGGACGACCATGAGCAGGGCCACCGAGGCCATCATTGACATCTGGGGCCGGGTCAGCACT
CCGGACCTAGATCCGGAGTCTATGGATCACGACTACGAATATGCCTTGGTCGACGGCAGGATCCTGATTCCGCGATTCCACTGGCAGACGATGTCGGGTGCATTTTCAGGGGCCGCGC
GCAAGGAGACAGATGATGTCGAGGTCGCTCTCAAGCACCTCAACATGTCGACTCCAGGTCTTCTACGCACAATGAAGTGGATTGACGGCAGCGTTCCGACCGCTCCCGCCAAGGGCGA
GGTGTTAGTCGAAGTGAAGGCCGTTGGTCTCAACTTCAGGGATGTTATTCTGGCTCTGGGCGTAGTCGAAGGGAATCCCAGCGGAATGGGCCACGAGGGCAGCGGTGTCATCCGCGCA
GTCGGACCAGATGTCCAAGACCTCTCCGTTGGCGACAGGGTCATGTTCATCCACGACGGCTGCTTCACCACGCAGCTGACGTTGTCGAAGGACATCTGCGTCAGGCTCGACGACTCGA
CGAGCTTTGTCCAGGGTGCCGCTCTGCCCGCCGTGCATGCAACCGCTCTGGCGGCACTGGTCGATGTTTCCAGGCTTCAAAGAGGACAGTCCGTGCTCATCCATGCCGCATGTGGTGG
CGTCGGTTTGGCCGCGATCCAGATTGCACAGAACATTGGTGCACAGATATACTGCACGGTAGGAAGCGAGCCGAAGCGGAGGTACCTCGAGGACACCTACAATATCCCGGCGGACCAC
ATCTTCAACTCCCGCGACGTGACGTTCCTTCCAGAGGTCATGCGCGCGACCAACGGACGCGGCGTTGACGTCGTCCTCAACTCGCTCTCCGGCGACCTCCTGCACGCGTCGTGGAAGT
GCGTCGCCGAGTTTGGCCTGATGGTAGAGATTGGGAAACGCGACTTCCAGCGCCGGTCAAAGCTCGCCATGGAGGTGTTCGAGGCGAACCGGAGCTTCGTGGGCCTGGACATACGCGG
GCTTTCCATCTCTCGCCCGGAGAGGGCCGCCGATCTGATGCGTCGATGCGTCGACATGATCCGGTCGAGGGCCATCCAGGGTCCTGTCGCGTGCACAACCTTCCCGGCCGTCGAGATC
CAGGATGCCTACAGGTACATGCAGAGCGCCAAGCATATCGGTAAGATTGTCATCGAGATGCCTGATGACCCGCGGGACCTTGGTGCAGGGGCAGCACCTCCAGAGGGCGAGTCGACCG
AGCTGGAAGTCTCTCCCAGGCCCAAGCCCAGCTTCCGCCCCGATCGTAGCTATCTCCTCGTGGGTGGTTTGGGAGGGCTGGGACGTGCTGTGGCAACCTGGATGGTGGAACACGGAGC
GCGGGTCTTGGTCTTCCTCTCCAGATCGGCCCGGGAGAATGATGACAACCGAGACTTCCTCAACGACCTCCGCTCCGAAGGTTGCACTGTTCTGTTGGTTCCCGGAAGCGTCAGCAAC
CTCGAGGATGTGCAAAAGGGGGTGGCAGTTGCTACGGCTGACCAGCCACTCGGCGGCCTCATCAACCTGTCCATGGTCCTCAGGGATGTCACGCTCAACAAGATGACATACTCGGACT
GGACCACGGCCGTGGAGCCAAAGGTCAGGGGAACGTGGAATCTGCATCACGCCACTGCAAACTGTACTTCCTTGGAGTTCTTCATACTGTTCTCTTCACAAAACGCTCAGATCGGCCA
ATGGGGCCAGGCCAACTACGCGGCTGCCAACACTTTCCTCGACGCCTTTGCCCAGTACCGGCACGGGCACAGCCTCGTCGCTTCGGTCATCGATGTCGGTCTCATGGGCGACGTGGGC
TTCGCGGCGGAGAACCGCGCGATCCTGAAGAAGCTGGGTAGGATCGGCATGTATATCCTGCAAGAGACAGACCTGCTCGATGCCATCAGCCTGGCTCTTCTGAAGTCACAGCCCATGC
ACGAGACGGAAGACAAGACGAGCCGCTACGTCACGCCGGGCTACGTGGGTATCGGCCTCAACACCACGACGCCCATGTCTGCCGCCTCGACCCGCGTCCCGTGGAAGCGAGACCCGCG
CATGAGCATCTACCACAACATGGATAACTCCAGCGGAGATGTCAGAGGCGAGGGCTCCTCCAAGGGCAGCTCGCTCAAGGTCGTCCTGGCCGCCGAGCCATCTGAGGAGAAGAAGACC
GAGATCATCGCCAAGGCCCTCGCCGGCACTCTCGGGAACTTCCTCATCAAAGACGGGAGCAGCTTCCCACTCGACAAGCCCCTCAAGATGCTCGGCATGGATTCTTTGATCGCCATGG
AGGTGCGGAACTGGATCCGACAGAACATCGGCGCCGAGACAAGCACGTTCACCGTGCTACAGAGCTCTTCCTTCATGCATCTGGCCGGCGAGATCAGGGCGGCGATGAATGCGGCGAG
CGAGGAGTGA
```

#### 3.2. Amino Acid Sequences

**pHis$_8$-SimG AT**

```
MKHHHHHHHH SSGLVPRGSHM-

        10         20         30         40         50         60
GNEKHADRGP NLLVFSAYSS ESLDRQISAY RDFAATHQGA SLKDLAYTLS SRRDHRPYRA

        70         80         90        100        110        120
YAIADDASGV QDASATVNAV VDGAPPPVGW IFTGQGAQWP EMGARLIDTS AAFRNRIRKL

       130        140        150        160        170        180
DKYLQTLKLE PPVSIEAELR KTKEDSRVHR PEMGHLVCVA MQVALVDVLR SWNIVPDFVL

       190        200        210        220        230        240
GHSSGETAAA YACGAITAEA AVYAATRRGI GNASSQRKGS MAAVGLGRDE IQPFLLEGVN

       250        260        270        280        290        300
IACENSQSNV TLSGDTEQVE SIVATLKAER PGVFARLLRV EKAYHSHHMC EYGPVYEEQL

       310        320        330        340        350        360
KPVVRSADPE IPFYSSVTGD RLTGEGQLGA SYWRANMENT VLFNPALRSA LRDRPGKLVL

       370        380        390        400        410        420
IELGPHPALE GPVGQILRDL GRTDDIYLGT CIRGSECDRS LLHLGGKLYQ QSIPMDLAAI

       430        440
CPPGAVLTNL PRYAWSQDTT HWEESR-
```

MW (including His-tag) = 51,161.72 Da

Catalytic Ser and His residues highlighted in Red.

**pHis$_8$-SimG_ACP**

```
MKHHHHHHHH SSGLVPRGSHM-

        10         20         30         40         50         60
GEGSSKGSSL KVVLAAEPSE EKKTEIIAKA LAGTLGNFLI KDGSSFPLDK PLKMLGMDSL

        70         80         90        100
IAMEVRNWIR QNIGAETSTF TVLQSSSFMH LAGEIRAAMN AASEE-
```

MW (including His-tag) = 13623.56 Da

Ppant attachment site highlighted Red.

**pHis$_8$-SimG KS-AT**

MKHHHHHHHH SSGLVPRGSHM-

10 20 30 40 50 60
APQQISPPNG TASPEAGTPR YTYRPASPAP ESPWADSAEG AELAKPMAII GMAMRLPGSV

70 80 90 100 110 120
RSGDDLWDLL STRKSGLCDI PKNRFNADGF YDPARGPGTI PVKKGYFLHD VQIEEFDTNV

130 140 150 160 170 180
FPIPKMELER LDPAQRQLLQ VAYECIENAG GTSWRGSRTG CYIGEFGEDF ADSSARESQQ

190 200 210 220 230 240
RGNLRYTGLA DFAIANRISY ELDLQGPSMV VKTACSSSLV CLDLACKAIQ SGECESALVG

250 260 270 280 290 300
GVSMLFSPAT YISVTDLGVI SPNGQCRSFD AGADGYARGE AVNMVMIKRL DHALRDKDPI

310 320 330 340 350 360
RAVIRGTGVN TDGRTNGMLT PSSAVQADLI RHTYEIAGID DLSQTAVVEC HGTGTPVGDP

370 380 390 400 410 420
IETEAVGRCF GDDGVTITSV KPNLGHAEAA AGLSSLIKCV LSLEHRQVLP NINFETPNPD

430 440 450 460 470 480
IPFEKYKLRV PTEVESWPQG RAERISINAF GIGGVNAHAI IESPAQFGIK KP**ANGIAGRG**

490 500 510 520 530 540
**VHTNGDSVPN GSQKQHEHDH RQDQSAYVKG DYMN**GTAVNG ANGNEKHADR GPNLLVFSAY

550 560 570 580 590 600
SSESLDRQIS AYRDFAATHQ GASLKDLAYT LSSRRDHRPY RAYAIADDAS GVQDASATVN

610 620 630 640 650 660
AVVDGAPPPV GWIFTGQGAQ WPEMGARLID TSAAFRNRIR KLDKYLQTLK LEPPVSIEAE

670 680 690 700 710 720
LRKTKEDSRV HRPEMGHLVC VAMQVALVDV LRSWNIVPDF VLGHSSGETA AAYACGAITA

730 740 750 760 770 780
EAAVYAATRR GIGNASSQRK GSMAAVGLGR DEIQPFLLEG VNIACENSQS NVTLSGDTEQ

790 800 810 820 830 840
VESIVATLKA ERPGVFARLL RVEKAYHSHH MCEYGPVYEE QLKPVVRSAD PEIPFYSSVT

850 860 870 880 890 900
GDRLTGEGQL GASYWRANME NTVLFNPALR SALRDRPGKL VLIELGPHPA LEGPVGQILR

910 920 930 940 950 960
DLGRTDDIYL GTCIRGSECD RSLLHLGGKL YQQSIPMDLA AICPPGAVLT NLPRYAWSQD

TTHWEESR

MW (including His-tag) = 106,963.24 Da (full length); 102,404.44 Da ($\Delta$473-514).

Catalytic Cys and His residues highlighted in Red.

Removed linker residues highlighted in Grey – gives rise to SimG KS-AT($\Delta$473-514).

#### 3.2. Supplementary Tables

**Table S1.** Primers for cloning of pHis$_8$-SimG.

| Construct | Forward Primer (5'-3') | Reverse Primer (5'-3') |
|---|---|---|
| pXW55-SimG | CCATCACCATCACCATCACCATCACACTAGTG CTCCTCAACAAATATCACCCCCCAATGG | CAAGCTCAGGACACACTTGATCAGGCTGG ACAG |
| | CTGTCACAAACCGCCGTCGTGGAGTGTC | GGTTGAAGAGAACCGTGTTCTCCATGTTCG |
| | GAGATCCCATTCTACTCCTCCGTCACCGGTG | CTGTCCAGAACGATGGCTTCGGGCTTCTG |
| | CATATCGAAGGATCCCAAGGAGCAGGGATTCA G | GATAATGAAAACTATAAATCGTGAAGGCAT TCACTCCTCGCTCGCCGCATTCATCGCCG C |

**Table S2.** NMR Data for 3R-hydroxyl-4R-methyl-6E-octenoic acid (Hma, **3**)

| | | |
|---|---|---|
| | 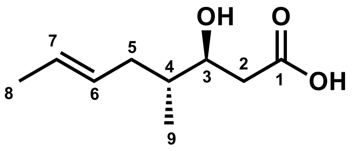<br>Data in agreement with reported literature.[2] | |
| Pos. | $\delta_C$ | $\delta_H$, multi, ($J$ in Hz) |
| 1 | - | |
| 2 | 38.2, $CH_2$ | Ha: 2.63 – 2.49, m, 1H<br>Hb: 2.49 – 2.37, m, 1H |
| 3 | 71.8, CH | 3.87, br, 1H |
| 4 | 38.7, CH | 1.66, overlap, 1H |
| 5 | 35.9, $CH_2$, | Ha: 2.21 – 2.13, m, 1H<br>Hb: 1.89, dt, $J$ = 14.3, 7.6, 7.6 Hz, 1H |
| 6 | 129.1, CH | 5.52 – 5.35, overlap, 1H |
| 7 | 127.2, CH | 5.52 – 5.35, overlap, 1H |
| 8 | 18.1, $CH_3$ | 1.66, d, $J$ = 5.6 Hz |
| 9 | 15.2, $CH_3$ | 0.88, d, $J$ = 6.6 Hz |

**Table S3.** Primers, annealing temperatures and restriction sites for cloning and mutagenesis of SimG KS-AT, AT and ACP constructs.

| Construct | Forward Primer (5'-3') | Reverse Primer (5'-3') | Temp. (°C) |
|---|---|---|---|
| **pHis$_8$-SimG KS-AT** | ATA**CATATG**GCTCCTCAACAAATATCA<br>(NdeI) | ATA**AAGCTT**TCACCTGGATTCTT<br>(HindIII) | 52 |
| **pHis$_8$-SimG AT** | ATA**CATATG**GGCAACGAAAAGCAT<br>(NdeI) | ATA**AAGCTT**TCACCTGGATTCTT<br>(HindIII) | 69 |
| **pHis$_8$-SimG ACP** | ATA**CATATG**GGCGAGGGCTCCTCCAA<br>(NdeI) | ATA**AAGCTT**TCACTCCTCGCTCGC CGCATT<br>(HindIII) | 69 |
| **Mutation** | **Forward Primer (5'-3')** | **Reverse Primer (5'-3')** | **Temp. (°C)** |
| **pHis$_8$-SimG KS-AT (Δ473-514)** | GGCACTGCCGTCAACGGG | AGGCTTCTTGATTCCAAACTGCG | 68 |
| **pHis$_8$-SimG KS-AT$^0$ (Δ473-514), (S705A)** | **AT$^0$**<br>ATAGGCAACGAAAAGCATGC<br>**pET28a(+)_KS**<br>ATAACGACGCACTGGGAAGAATCCA | **AT$^0$**<br>ATATCACCTGGATTCTTCCCA<br>**pET28a(+)_KS**<br>ATATCCGCGGTCAGCATGCTTTT | 70 |
| **pHis$_8$-SimG KS$^0$-AT$^0$ (Δ473-514), (C215A, S705A)** | CAAGACGGCTGCATCGTCTTCCCTCGTG TG | ACCACCATACTCGGGCCT | 65 |
